# Identification of Fasnall as a therapeutically effective Complex I inhibitor

**DOI:** 10.1101/2024.05.03.592013

**Authors:** Dzmitry Mukha, Jena Dessain, Seamus O’Connor, Katherine Pniewski, Fabrizio Bertolazzi, Jeet Patel, Mary Mullins, Zachary T. Schug

## Abstract

Proliferating cancer cells actively utilize anabolic processes for biomass production, including *de novo* biosynthesis of amino acids, nucleotides, and fatty acids. The key enzyme of the fatty acid biosynthesis pathway, fatty acid synthase (FASN), is widely recognized as a promising therapeutic target in cancer and other health conditions^1,2^. Here, we establish a metabolic signature of FASN inhibition using a panel of pharmacological inhibitors (GSK2194069, TVB-2640, TVB-3166, C75, cerulenin, and Fasnall). We find that the activity of commonly used FASN inhibitors is inconsistent with the metabolic signature of FASN inhibition (accumulation of malonate, succinate, malonyl coenzyme A, succinyl coenzyme A, and other metabolic perturbations). Moreover, we show that one of these putative FASN inhibitors, Fasnall, is a respiratory Complex I inhibitor that mimics FASN inhibition through NADH accumulation and consequent depletion of the tricarboxylic acid cycle metabolites. We demonstrate that Fasnall impairs tumor growth in several oxidative phosphorylation-dependent cancer models, including combination therapy-resistant melanoma patient-derived xenografts. Fasnall administration does not reproduce neurological side effects in mice reported for other Complex I inhibitors^3,4^. Our results have significant implications for understanding the FASN role in human health and disease and provide evidence of therapeutic potential for Complex I inhibitors with fast systemic clearance. Our findings also highlight the continuing need for validation of small molecule inhibitors to distinguish high-quality chemical probes and to expand the understanding of their application.

---

Fatty acid biosynthesis integrates multiple resources, including carbon, energy, and reducing equivalents supplied by several major parts of central metabolism. Fatty acid biosynthesis was shown to be a crucial anabolic process for cancer tumor growth and tissue invasion^1,2^. Breast cancer brain metastases experience low lipid availability and depend on *de novo* fatty acid biosynthesis for growth^5^. The upregulation of FASN promotes cancer cell motility and metastasis^6^. Newly synthesized fatty acids modulate cellular levels of lipid saturation and promote cancer cell resistance to lipid peroxidation and ferroptosis^7–9^.

The final steps of *de novo* fatty acid biosynthesis are catalyzed by fatty acid synthase (FASN; Fig. 1a). Over the last three decades, many pharmacological agents were developed to inhibit FASN activity in cells, such as C75^10^, Orlistat^11^, BI 99179^12^, GSK2194069^13^, TVB-2640^14,15^ (Denifanstat, ASC40), TVB-3166^16^, Fasnall^17^, among others^1^. FASN inhibitors, along with the genetic manipulation of gene expression, are the major tools for studying the role of FASN in various contexts. By the end of 2023, at least seven clinical trials dedicated to FASN inhibition efficacy against multiple cancers were registered (Supplementary Table 1).

**Fig. 1.**
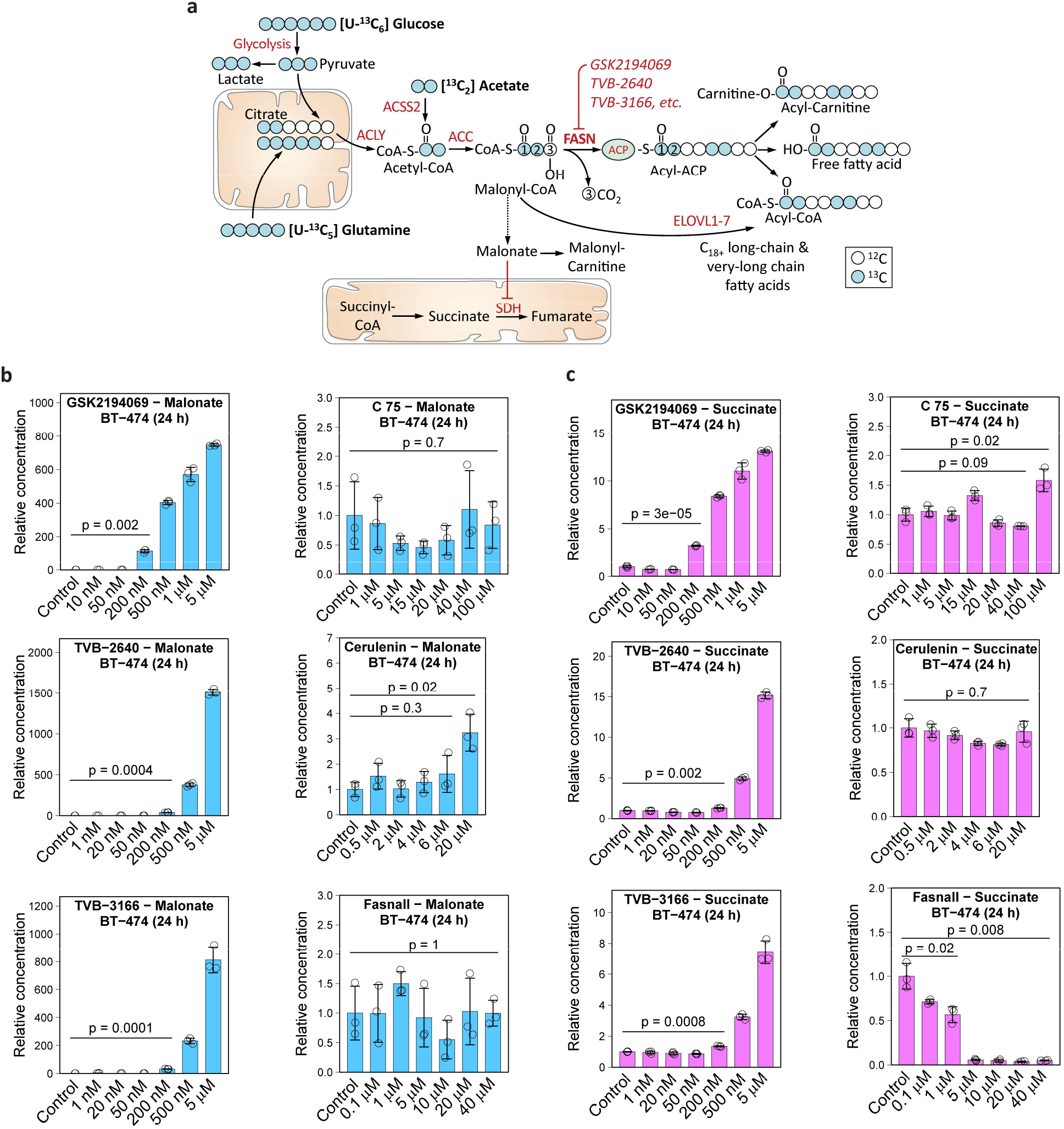
Malonate and succinate accumulate in response to three FASN inhibitors, GSK2194069, TVB-2640, and TVB-3166, but not C75, cerulenin, and Fasnall. a, Schematic of the *de novo* fatty acid biosynthesis pathway showing entry points of carbon originating from [U-^13^C_6_] D-glucose, [U-^13^C_5_] L-glutamine, and [^13^C_2_] acetate. b-c, Relative concentrations of malonate (b) and succinate (c) in BT-474 cells treated with six compounds previously described as FASN inhibitors. Data are mean ± SD (n = 3). The relative concentrations of malonyl-CoA, succinyl-CoA, and acetyl-CoA are provided in Extended Data Fig. 1.

Acetyl coenzyme A (acetyl-CoA) is the carbon source for *de novo* fatty acid biosynthesis. In most cells, acetyl-CoA is primarily produced from citrate, and therefore, fatty acid biosynthesis has an intricate connection to the tricarboxylic acid (TCA) cycle performance. One of the main functions of the TCA is NADH oxidation to NAD^+^. *De novo* fatty acid biosynthesis also consumes reducing equivalents, but in the form of NADPH. Surprisingly, *de novo* fatty acid biosynthesis exhibits a paradoxical dependency on NAD^+^ produced by the functional TCA cycle^18^. Moreover, the expression of lipid biosynthesis genes is significantly anti-correlated with hypoxia-induced markers in tumors^18^.

Considering the interconnection of fatty acid biosynthesis and the TCA cycle, we hypothesize that FASN inhibition might have a profound impact on cell metabolism by affecting metabolites upstream of the FASN. In this work, we explore the metabolic perturbations and create a robust biomarker profile of FASN inhibition. Using the metabolic signature of polar metabolite markers, we conclude that several previously described FASN-targeting pharmacological agents do not inhibit FASN in cells. Accordingly, we suggest revising the existing body of knowledge on the mechanistic role of FASN in various human health conditions.

## Results

### Metabolic signature of fatty acid synthase inhibition

Based on the known structure of the fatty acid biosynthesis pathway (Fig. 1a), we proposed and tested several predictions about the effect of FASN inhibition on cell metabolism. First, we expect the accumulation of the FASN substrate, malonyl-coenzyme A (malonyl-CoA). Hydrolysis of malonyl-CoA, whether spontaneous or mediated by a yet-undescribed enzyme, can release free malonate, which, in turn, acts as an inhibitor of the mitochondrial TCA cycle enzyme succinate dehydrogenase (SDH)^19^. Inhibition of SDH should manifest in the accumulation of succinate and succinyl-CoA (Fig. 1a).

We analyzed the effect of six widely used pharmacological FASN inhibitors on the metabolome of BT-474 breast cancer cells (Extended Data Fig. 1a). As we show below, only three of the compounds – GSK2194069, TVB-2640, and TVB-3166 – demonstrate a consensus metabolic signature expected for FASN inhibition, while three other compounds – C75, cerulenin, and Fasnall – predominantly act on other molecular targets in cells. Three consensus drugs cause the accumulation of intracellular malonate (750 to 1500-fold) and succinate (7 to 15-fold) in a dose-dependent manner (Fig. 1b-c). The increase of malonyl-CoA concentration was in the range of 15 to 30-fold (Extended Data Fig. 1b). Succinyl-CoA and acetyl-CoA were less affected by FASN inhibition, with only a minor 1.5- to 2-fold accumulation of the latter (Extended Data Fig. 1c).

Using a mass spectrometry-based approach, previous studies showed that GSK2194069, BI 99179, and orlistat (Extended Data Fig. 1e) caused malonyl-CoA accumulation in cells, corroborating our results for GSK2194069 and further expanding the set of compounds with the consensus metabolic signature^13,20^. The accumulation of O-malonylcarnitine, another malonate metabolism product, was shown in patients receiving TVB-2640 treatment^21^, similar to our results *in vitro* in BT-474 cells (Extended Data Fig. 2a). We confirmed the ability of malonate and its methyl diester, dimethylmalonate (DMM), to cause the intracellular accumulation of malonate and succinate (Extended Data Fig. 2b). Malonyl-CoA concentration is unaffected by DMM (Extended Data Fig. 2b). The stereotypical effect of DMM treatment was confirmed in seven other breast cancer cell lines (Extended Data Fig. 2c-d). The co-treatment of cells with DMM and GSK2194069 combines the effects of both compounds and leads to malonyl-CoA accumulation (Extended Data Fig. 2e).

Using liquid chromatography-tandem mass spectrometry (LC-MS/MS), we confirmed the intracellular presence of five compounds (GSK2194069, TVB-2640, TVB-3166, C75, and Fasnall) in BT-474 cells (Extended Data Fig. 2f). One remaining compound from our panel, cerulenin^22^, was found lethal at 50 µM for BT-474 cells in a 24-h experiment, confirming its uptake by cells. C75, cerulenin, and Fasnall do not cause the expected accumulation of malonate, succinate, malonyl-CoA, succinyl-CoA, and acetyl-CoA. Instead, Fasnall treatment leads to a dose-dependent depletion of succinate, succinyl-CoA, malonyl-CoA, and acetyl-CoA (Fig. 1b-c, Extended Data Fig. 1b-d).

### Total metabolome profiling reveals the lack of FASN inhibition signature in cells treated with C75, cerulenin, and Fasnall

We analyzed the relative changes in intracellular and extracellular metabolomes for thirty conditions corresponding to different drugs and dosages. Unsupervised clustering analysis revealed deviation of C75, cerulenin, and Fasnall from the other three FASN inhibitors (Fig. 2a). Surprisingly, high concentrations of Fasnall lead to metabolic perturbations that are not reproduced by any other drug in our FASN inhibitor panel. Considering that Fasnall has been previously reported to exhibit anti-cancer activity *in vivo*^17^, we next focused on comparing Fasnall with one representative compound from the consensus group, GSK2194069. To validate the chemical structure of both compounds and establish a molecular fingerprint for every commercially purchased compound batch, we recorded a collision-induced dissociation (CID) profile for five major molecular fragments of both drugs (Extended Data Fig. 3a-d). Metabolites affected by GSK2194069 are associated with the biosynthesis and turnover of fatty acids (Fig. 2b). However, Fasnall at 1 µM (∼3.7 times below the reported IC_50_ concentration^17^) triggers significant perturbations mainly in glycolysis, redox balance, and the TCA cycle (Fig. 2c; Supplementary Tables 2-4). We assessed whether metabolic perturbations induced by Fasnall can bias resazurin reduction into resorufin, a commonly used cell proliferation assay. Indeed, a short 1.5 h exposure to Fasnall leads to a significant decrease in resorufin fluorescence in a panel of eight breast cancer cell lines, while GSK2194069 has no significant effect (Fig. 2d-e, Extended Data Fig. 3e-f). The data suggest that Fasnall quickly rewires cell metabolism by affecting reactions beyond *de novo* fatty acid biosynthesis.

**Fig. 2.**
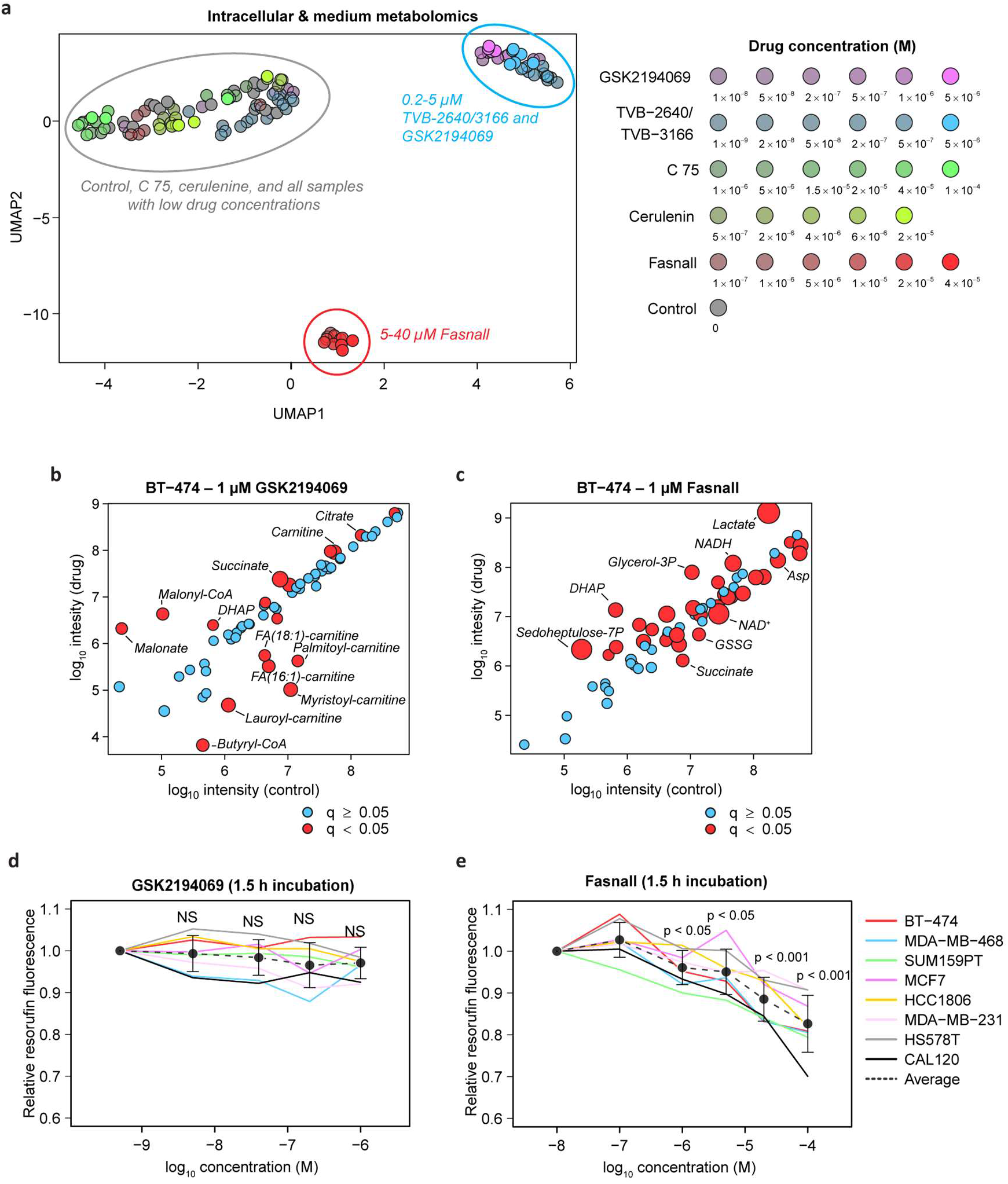
GSK2194069, TVB-2640, and TVB-3166 form a consensus metabolic profile of FASN inhibition. a, UMAP 2D projection of 208-mer vectors containing relative changes of intracellular and medium metabolite concentrations in response to the corresponding drug treatment of BT-474 cells. 123 dots represent 266 LC-MS samples in total, one intracellular and one medium sample per Petri dish. b-c, Perturbations in intracellular metabolite concentrations in BT-474 cells treated with 1 μM GSK2194069 (b) and 1 μM Fasnall (c) for 24 hours. Log10 integrated LC-MS peak intensity is used for both axes. Metabolites with Benjamini-Hochberg FDR-adjusted p-values (q-values) lower than 0.05 are depicted with red circles. The size of the red circles is inversely proportional to the q-values. d-e, Relative resorufin fluorescence in eight breast cancer cell lines treated with different concentrations of GSK2194069 (d) and Fasnall (e) for 1.5 h. Data on panels d and e are mean ± SE (n ≥ 12).

### Fasnall, unlike FASN inhibitors, decreases cancer cell proliferation in standard culture conditions

To further test if the anti-cancer activity of Fasnall can be explained by FASN inhibition, we conducted a cell proliferation assay. The antiproliferative effects of FASN inhibitors have been previously demonstrated in special culture conditions with decreased content or complete lack of lipids from fetal bovine serum (FBS)^13,16,23^. GSK2194069 did not affect the proliferation of breast cancer cells in regular, lipid-containing culture conditions (Fig. 3a-b). In contrast, Fasnall decreased cancer cell proliferation, even in the presence of serum lipids (Fig. 3c). Considering a previous observation of a negative correlation between lipid biosynthesis and tumor hypoxia expression markers in primary tumor samples^18^, we tested whether the sensitivity to Fasnall is correlated to the sensitivity to hypoxia. Indeed, we find that the effect of Fasnall has a significant correlation with sensitivity to 1% O_2_ (Fig. 3e). In sum, lipid availability does not mitigate Fasnall cell toxicity, unlike the FASN inhibitor GSK2194069. We show a significant correlation between sensitivity to Fasnall and growth inhibition by hypoxia. It was previously reported that the latter is correlated with lipid biosynthesis activity in cancer^18^. However, our data argue against a causal link between sensitivity to Fasnall and lipid biosynthesis inhibition.

**Fig. 3.**
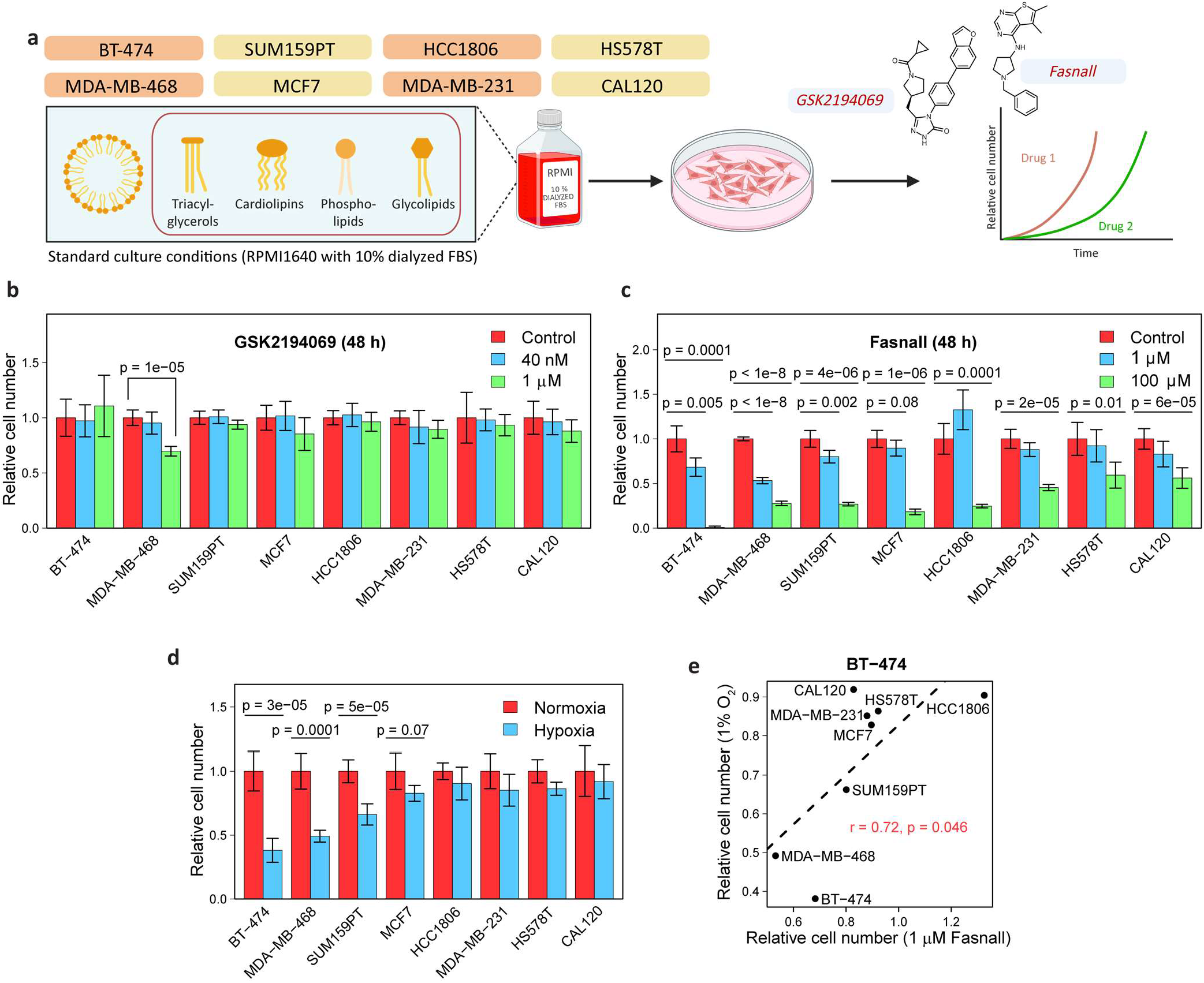
Fasnall inhibits cancer cell proliferation in presence of lipids in the medium. **a**, Schematic of the cell proliferation experiment. FASN inhibition was applied in the presence of lipids from fetal bovine serum (FBS). **b-c**, Cell number (relative to control) for eight cancer cell lines treated with GSK2194069 (**b**) and Fasnall (**c**) for 48 h. **d**, Cell number after 48 h growth in hypoxia (relative to normoxia). **e**, Comparison of the effect of Fasnall vs. hypoxia on cell proliferation relative to growth in regular culture conditions. Data are mean ± SD (n = 3).

### Fasnall treatment decreases fatty acid synthesis from glucose but not glutamine

To dissect the mode of action of Fasnall, we decided to trace the utilization of glucose and glutamine, two primary carbon sources for fatty acid biosynthesis in cultured cancer cells (Fig. 1a; most cell culture media are not supplemented with acetate). Carbon is incorporated in fatty acids in two-carbon units, with even-numbered isotopologues getting labeled with ^13^C in isotope tracing experiments (Fig. 1a). The main product of FASN is palmitic acid (16:0), while longer acyl chains are generated via elongation in endoplasmic reticulum^2^. In BT-474 cells, GSK2194069 virtually eliminates ^13^C labeling from [U-^13^C_6_] D-glucose and [U-^13^C_5_] L-glutamine in myristic (14:0), palmitic, and palmitoleic (16:1) acids (Fig. 4a). Stearic (18:0) and oleic (18:1) acids demonstrate inclusion of a single two-carbon fragment via the elongation of the acyl chain catalyzed by elongases (Fig. 4a; Extended Data Fig. 4a). Fasnall treatment decreases glucose contribution to fatty acid labeling in cells fed with [U-^13^C_6_] D-glucose (Fig. 4a), corroborating previous results obtained with [^3^H] D-glucose tracing^17^. Remarkably, glutamine contribution to carbon labeling in fatty acids significantly increases and fully compensates for the loss of glucose-derived flux in myristic and palmitic acids. Moreover, GSK2194069 suppresses the abundance of medium- and long-chain O-acylcarnitines, while Fasnall fails to do so (Fig. 4b). In sum, isotope glucose tracing is not sufficient to support fatty acid biosynthesis inhibition by Fasnall. Also, the data do not indicate that Fasnall can perturb the relative concentrations of fatty acids bound to carnitine.

**Fig. 4.**
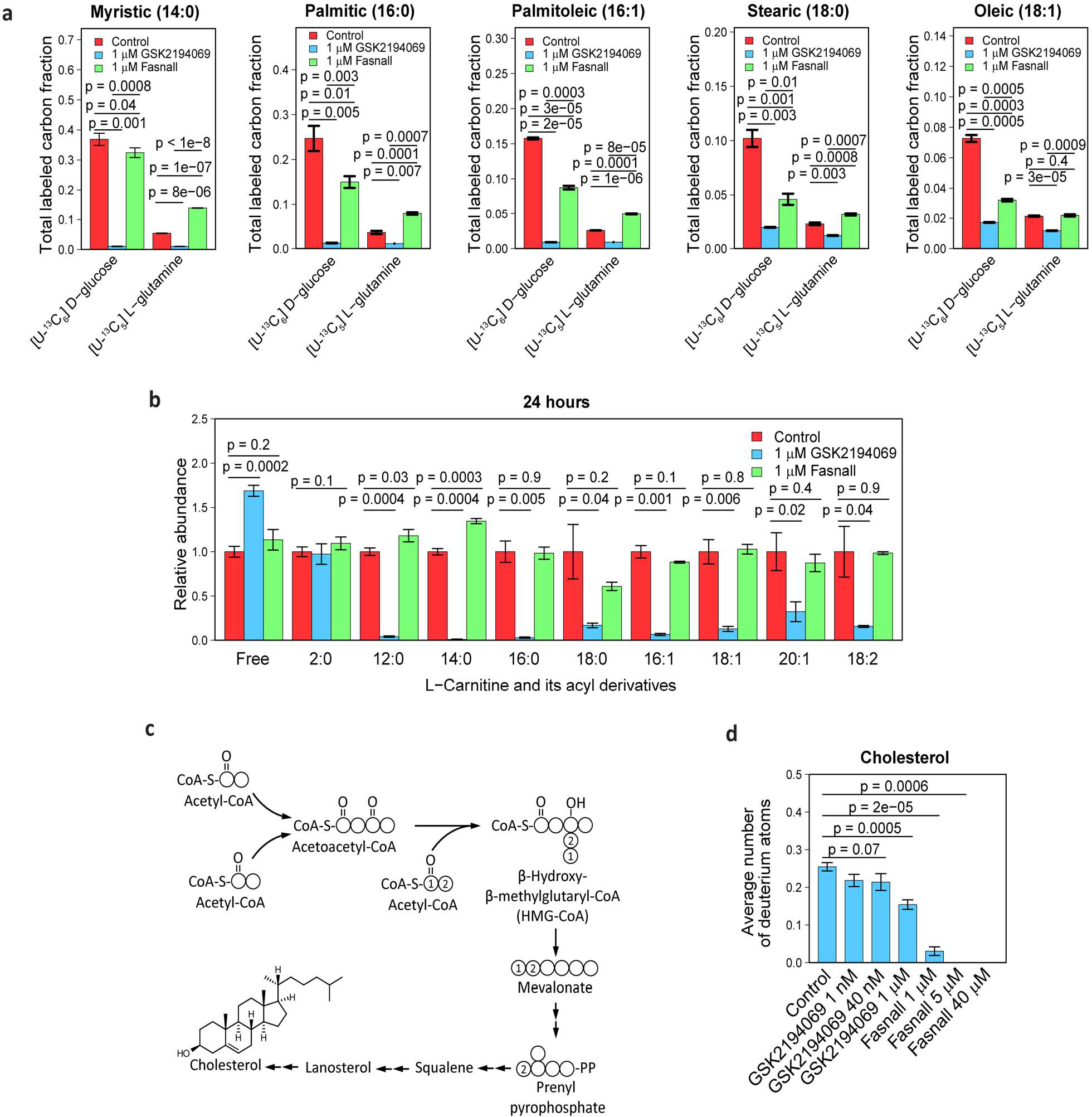
Isotope tracing demonstrates that Fasnall does not target FASN. **a**, Total ^13^C labeling of (left-to-right) myristic, palmitic, palmitoleic, stearic, and oleic acids in BT-474 cells fed with [U-^13^C_6_] D-glucose and [U-^13^C_5_] L-glutamine for 24 h. The decrease in the relative contribution of glucose to fatty acid biosynthesis in Fasnall-treated cells is compensated by glutamine. **b**, Fasnall does not perturb the abundance of O-acyl-carnitines in BT-474 cells. **c**, Schematic of cholesterol biosynthesis. **d**, Fasnall inhibits cholesterol biosynthesis, as evident from the deuterium labeling of cholesterol in BT-474 cells cultured in RPMI-1640 medium with 20% D_2_O for 24 h. Data are mean ± SD, n = 3.

### Fasnall inhibits cholesterol biosynthesis and the elongation of fatty acids

The mevalonate pathway fuels cholesterol biosynthesis without involving malonyl-CoA and instead using acetoacetyl coenzyme A and β-hydroxy-β-methylglutaryl coenzyme A as early precursors (Fig. 4c). Isotopic tracing in RPMI-1640 medium containing 20% v/v deuterium oxide (D_2_O) revealed that Fasnall dramatically decreases the ability of cells for *de novo* cholesterol biosynthesis in BT-474 at 1 µM, and completely ablates it at 5 µM (Fig. 4d, Extended Fig. 5). Deuterium labeling of myristic acid remains unaffected at 1 µM Fasnall and remains readily detectable even at 40 µM (∼7.5% of the control level at the drug concentration ∼10-fold higher than the reported IC_50_). In contrast, ximenic acid (26:1) labeling, a readout for elongase activity, loses all labeling at 5 µM Fasnall (Extended Fig. 5b-e). GSK2194069 has a lower impact on cholesterol biosynthesis and does not affect elongase activity, while deuterium labeling in myristic acid is completely undetectable at 1 µM GSK2194069 (Extended Fig. 5). The isotope tracing data demonstrate that the Fasnall treatment impacts multiple pathways that use acetyl-CoA as an intermediate. However, the perturbations in fatty acid carbon labeling cannot be explained by FASN inhibition.

**Fig. 5.**
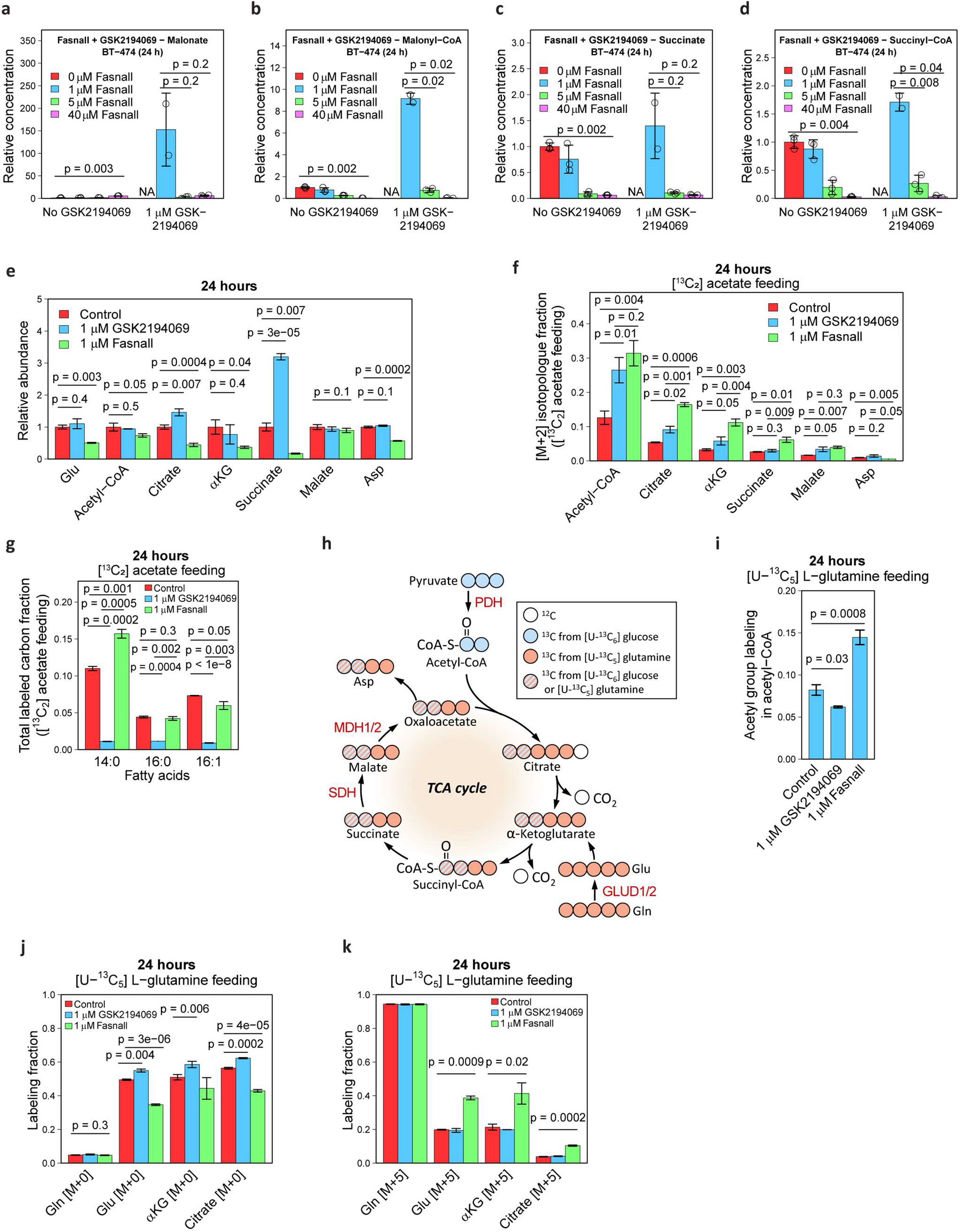
Fasnall acts upstream of FASN, affecting the TCA cycle and activating reductive carboxylation. **a-d**, Relative concentrations of malonate (**a**), malonyl-CoA (**b**), succinate (**c**), and succinyl-CoA (**d**) in BT-474 cells treated with a combination of 1 µM GSK2194069 and various concentrations of Fasnall for 24 h. **e**, Relative abundance of the TCA cycle metabolites and aspartate in BT-474 cells treated with 1 µM GSK2194069 and 1 µM Fasnall for 24 h. **f-g**, Fraction of [M+2] isotopologue in the TCA cycle metabolites and aspartate (**f**), as well as the total ^13^C labeling in myristic, palmitic, and palmitoleic acids (**g**) in BT-474 cells fed with 200 µM [^13^C_2_] acetate. **h**, Schematic of [U-^13^C_6_] D-glucose and [U-^13^C_5_] L-glutamine tracing in the TCA cycle. **i**, ^13^C labeling in the acetyl group of acetyl-CoA in BT-474 cells fed with [U-^13^C_5_] L-glutamine. **j-k**, Fraction of [M+0] (**j**) and [M+5] (**k**) isotopologues in glutamine, glutamate, α-ketoglutarate, and citrate in cells fed with [U-^13^C_5_] L-glutamine. Data are mean ± SD, n = 3.

### Fasnall acts upstream of FASN

Next, we assessed the order of enzymatic targets of Fasnall and GSK2194069 by co-treating BT-474 cells with both drugs. The Fasnall-specific metabolic response, characterized by a lack of malonate accumulation and decreased malonyl-CoA, succinate, and succinyl-CoA concentrations, prevails in cells co-treated with both compounds (Fig. 5a-d). Increasing the Fasnall concentration to 40 µM does not revert this trend. This indicates that Fasnall likely acts on a target upstream of FASN and that this effect is the primary consequence of Fasnall treatment until the concentration of Fasnall becomes lethal to cultured cancer cells. In support of this premise, we detect a decrease in concentrations of multiple TCA cycle metabolites, as well as aspartate in cells treated with 1 µM Fasnall (Fig. 5e). Next, we used 200 µM [^13^C_2_] acetate supplementation to provide a carbon source that can bypass pyruvate dehydrogenase (PDH) in mitochondria and ATP citrate lyase (ACLY) in the cytosol of BT-474 cells by generating acetyl-CoA via acetyl-CoA synthetases ACSS1 and ACSS2, respectively^24^ (Fig. 1a). [^13^C_2_] Acetate-fed cells have a significantly higher fraction of [M+2] isotopologues in the TCA cycle metabolites in the presence of 1 µM Fasnall than control and GSK2194069-treated cells, indicating a decreased contribution of glucose to the TCA cycle (Fig. 5f). Myristic acid labeling in [^13^C_2_] acetate-fed cells treated with Fasnall is higher than in control cells, while palmitic and palmitoleic acids display labeling pattern that matches the control cells. GSK2194069 at 1 µM concentration effectively prevents ^13^C incorporation in C_14_-C_16_ fatty acids in cells fed with [^13^C_2_] acetate, supporting FASN inhibition by GSK2194069 (Fig. 5g, Extended Data Fig. 6a-c). Thus, bypassing PDH allows cells to use acetate carbon to maintain FASN activity in the presence of Fasnall.

### Fasnall increases TCA cycle anaplerosis from glucose and lowers PDH flux

Our data prompted us to assess the effects of Fasnall on the TCA cycle activity and acetyl-CoA generation. The ^13^C labeling of the acetyl group in acetyl-CoA confirms the increased relative contribution of glutamine (Fig. 5h-i, Extended Data Fig. 6d-e). Citrate [M+5], produced via reductive carboxylation in the TCA cycle in cells fed with [U-^13^C_5_] L-glutamine, is further used for generating acetyl-CoA in the cytosol via ACLY (Fig. 1a, 5h). The remaining carbon re-enters the TCA cycle as malate [M+3], being further converted into oxaloacetate and aspartate (Fig. 5j-k, Extended Data Fig. 6f-h). The activation of reductive carboxylation supports the observation of decreased flux from glucose and suggests the accumulation of reducing equivalents in mitochondria.

[U-^13^C_6_] D-Glucose tracing in BT-474 cells treated with Fasnall demonstrates a dose-dependent departure from [M+2] malate labeling, derived from the PDH flux, towards [M+3] labeling from anaplerosis involving malic enzymes (ME1/2/3) and/or pyruvate carboxylase (PC, Fig. 6a-b, Extended Data Fig. 7a). Additionally, cells accumulate reduced cofactor NADH and glycerol 3-phosphate generated by reduction of dihydroxyacetone phosphate (DHAP) using an electron from NADH. These changes occur as early as 4 h after Fasnall treatment (Fig. 6c-d) and can be observed after 24 h (Fig. 6e-f). Fasnall-treated cells become highly glycolytic and increase their glucose consumption flux ∼9-fold and their lactate secretion flux ∼18-fold, indicating that virtually all extra glucose consumption flux is diverted to lactate secretion according to the 1:2 stoichiometry (Fig. 6g-h; Extended Data Fig. 7b). Both lactate and pyruvate are also accumulated intracellularly (Fig. 6i), while ATP and the TCA cycle metabolite pools are depleted (Fig. 6j-l). In contrast, FASN inhibitor GSK2194069 does not affect glucose consumption and lactate secretion (Fig. 6g-l).

**Fig. 6.**
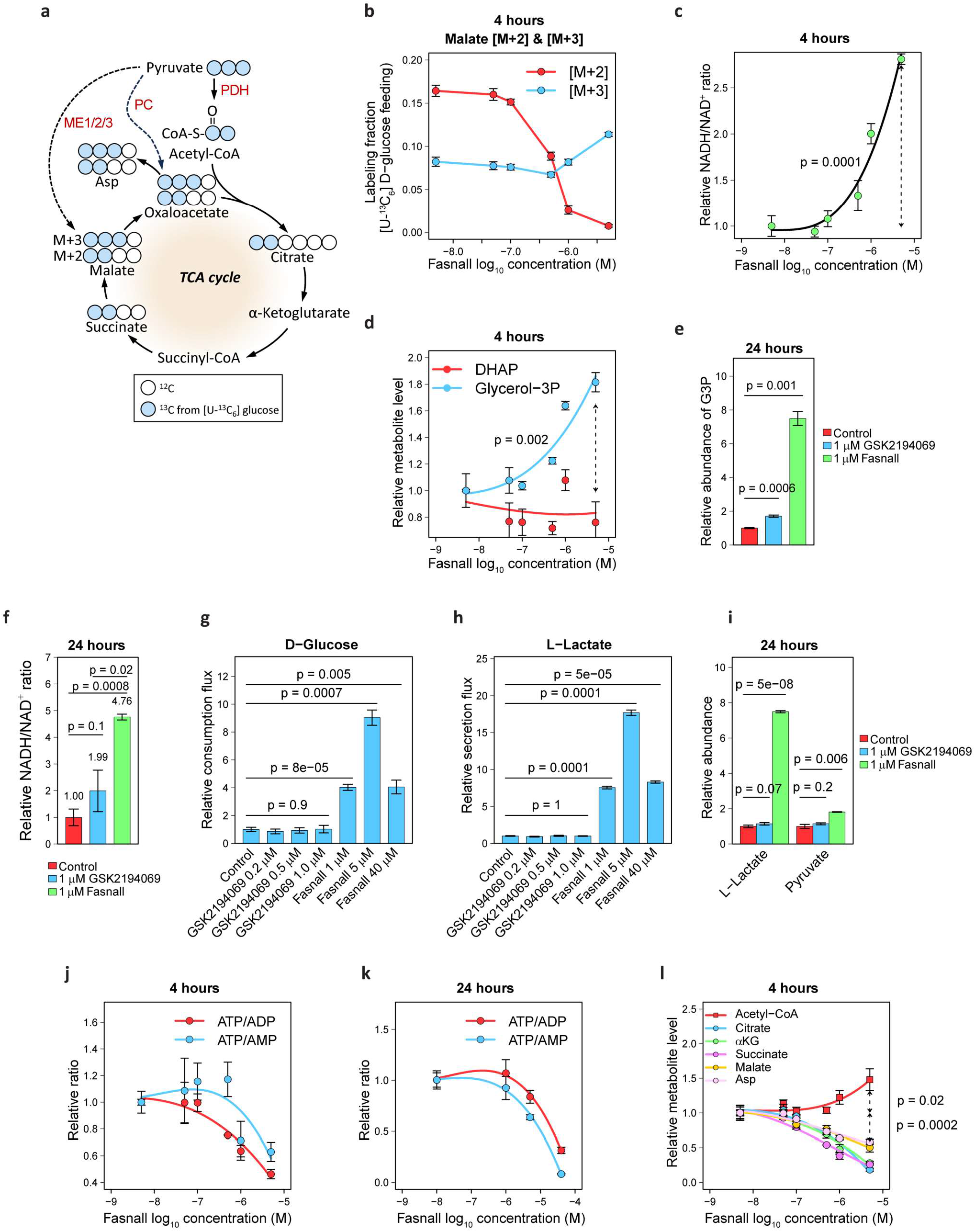
Perturbations in central carbon metabolism caused by Fasnall. **a**, Schematic of ^13^C carbon tracing in the TCA cycle in cells fed with [U-^13^C_6_] D-glucose. **b**, Switch from the predominant M+2 to M+3 malate labeling in OMM1.3 cells treated with Fasnall for 4 h. **c-f**, Concentration-dependent increase in the NADH/NAD^+^ ratio (**c**) and glycerol 3-phosphate concentration (**d**) in OMM1.3 cells treated with GSK2194069 and Fasnall for 4 h and 24 h (**e-f**). **g-i**, Glucose consumption (**g**) and lactate secretion flux (**h**), as well as lactate and pyruvate intracellular concentrations in BT-474 cells treated with GSK2194069 and Fasnall for 24 h. **j-k**, ATP/ADP and ATP/AMP ratios in OMM1.3 (**j**) and BT-474 (**k**) cells treated with Fasnall and for 4 h and 24 h, correspondingly. **l**, TCA cycle metabolite depletion caused by Fasnall treatment in OMM1.3 cells. Data are mean ± SD (n = 3).

### Fasnall does not target PDH or malate dehydrogenase (MDH)

The glycolytic switch caused by Fasnall may have multiple competing explanations, all pointing towards dysfunction of the TCA cycle, pyruvate utilization in mitochondria, and/or NADH oxidation. Blockade of PDH may prevent pyruvate oxidation in mitochondria and force its reduction to lactate. Alternatively, malate dehydrogenase (MDH) inhibition might disrupt the malate-aspartate electron shuttle, which maintains redox balance in the cytosol.

We ruled out a direct Fasnall inhibition of PDH in an assay utilizing ammonium-sulfate-precipitated and reconstituted total protein from BT-474 cells, with purified PDH as a positive control (Fig. 7a, Extended Data Fig. 8a). Also, Fasnall does not inhibit MDH from porcine heart in the corresponding reconstituted activity assay (Fig. 7b, Extended Data Fig. 8b). Moreover, LW6, an MDH2 inhibitor^25^, causes ∼40% increase in malate concentration – an effect that we do not observe in cells treated with Fasnall (Extended Data Fig. 8c). The data support the possibility that Fasnall inhibits Complex I.

**Fig. 7.**
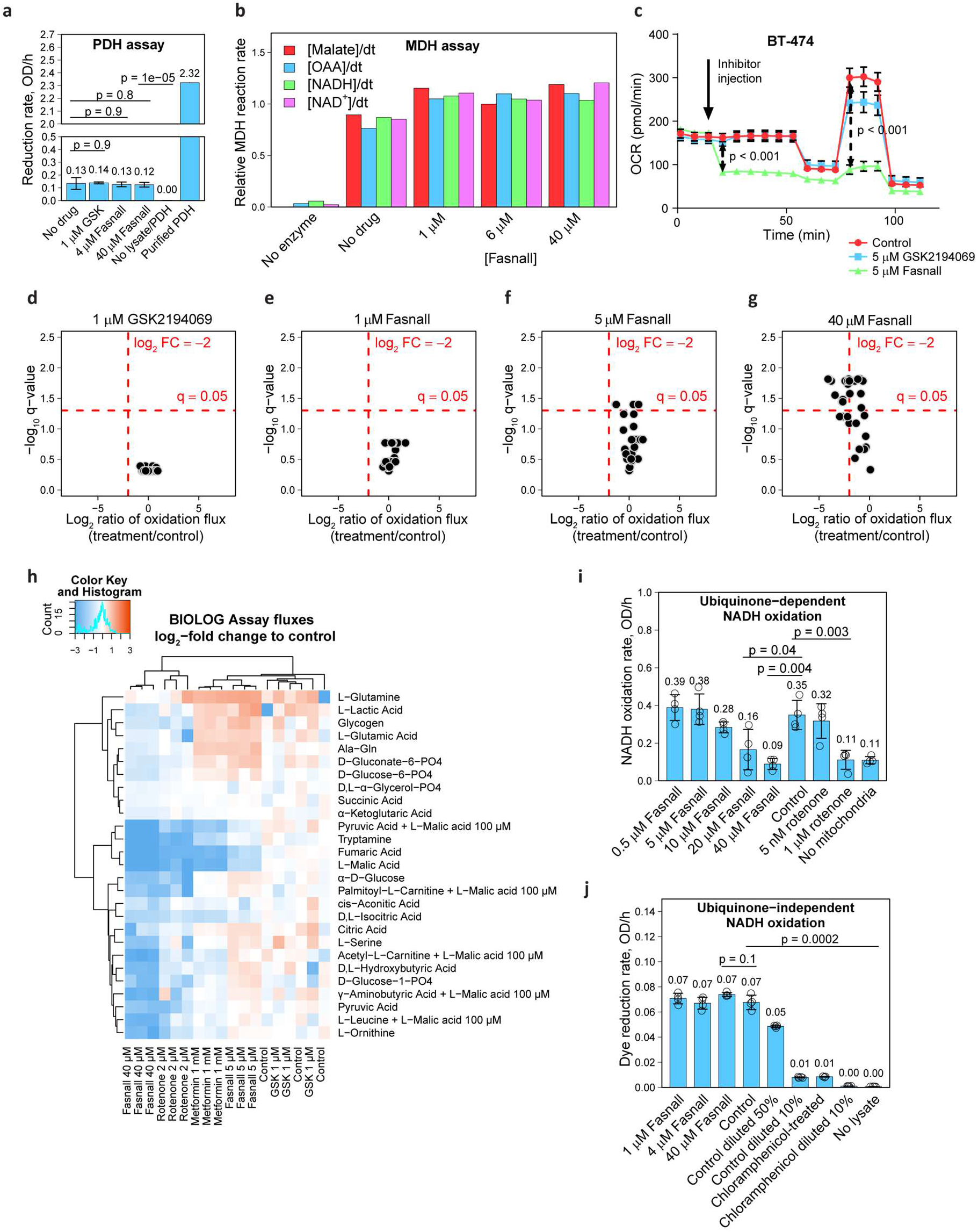
Fasnall does not inhibit PDH or MDH *in vitro*; instead, it impairs the oxidation of various metabolites, mimicking rotenone and metformin. a-b, PHD (**a**) and MDH (**b**) in vitro assays**. c**, Oxygen consumption assay with injecting with 5µ GSK2194069 and Fasnall. **d-h**, Selected volcano plots (**d-g**) and relative changes of mitochondrial substrate oxidation fluxes (**h**) in the BIOLOG assay. **i-j**, Complex I assays with ubiquinone (**i**) and a calorimetric reporter dye (**j**) as the final electron acceptor. Data on panel **a, i-j** are mean ± SD (n = 4).

### Fasnall acts similar to known Complex I inhibitors and prevents mitochondria from oxidizing substrates

The addition of Fasnall in the Seahorse assay causes an immediate and significant decrease in oxygen consumption (Fig. 7c). The GSK2194069 effect is limited to the decline in the maximal respiration capacity, which hints at the potential manifestation of SDH inhibition caused by malonate (Fig. 7c). To rule out a regulatory influence of Fasnall on glycolysis, we performed a BIOLOG phenotypic assay on permeabilized BT-474 cells, measuring their capacity for oxidizing 29 different substrates. As expected, GSK2194069 does not alter oxidation flux in the absence of malonate accumulation (Fig. 7d). Fasnall, in contrast, demonstrates a dose-dependent decrease in the reporter dye reduction, with the most potent effect against malate, fumarate, and hydroxybutyrate. All three metabolites are oxidized in reactions generating NADH and mainly rely on respiratory Complex I for NAD^+^ recycling. We extended the assay to include rotenone and metformin to validate that the observed phenotypic pattern is reminiscent of Complex I inhibition. Fasnall, rotenone, and metformin formed a cluster of highly correlated responses (Fig. 7h). A cell-free ubiquinone-dependent NADH oxidation assay demonstrates dose-dependent inhibition of Complex I activity by Fasnall (Fig. 7i). In a ubiquinone-independent assay, with a dye serving as a terminal electron acceptor, Fasnall does not impact the rate of NADH oxidation (Fig. 7j). The combined analysis of metabolite extracts from cells and medium, analogous to the one presented in Fig. 2a and extended to include rotenone and LW6, confirmed that treatment with Fasnall and rotenone induces highly similar metabolic changes (Extended Data Fig. 8d; Pearson’s correlation of 2 µM rotenone to 20 µM Fasnall *r* ∈ (0.54 − 0.58), *p* < 5 × 10^-9^). Thus, our data suggest that Fasnall is likely a ubiquinone-dependent Complex I inhibitor.

### Zebrafish embryos accumulate lactate upon Fasnall treatment

Rotenone is known for its toxicity in fish. To compare the metabolic effects and toxicity of Fasnall with rotenone *in vivo*, we exposed zebrafish embryos 48 h post-fertilization to a drug-containing medium for 6 h (Fig. 8a). Rotenone at 25 nM is lethal to fish embryos, while 5 nM concentration leads to a ∼15-fold lactate accumulation. Similarly, Fasnall treatment increases lactate content, although the magnitude of the effect is significantly lower (Fig. 8b). Unlike 5 nM rotenone, Fasnall treatment does not cause visible phenotypic changes in the yolk (Extended Data Fig. 9a). The zebrafish model suggests that Fasnall acts as a Complex I inhibitor *in vivo*.

**Fig. 8.**
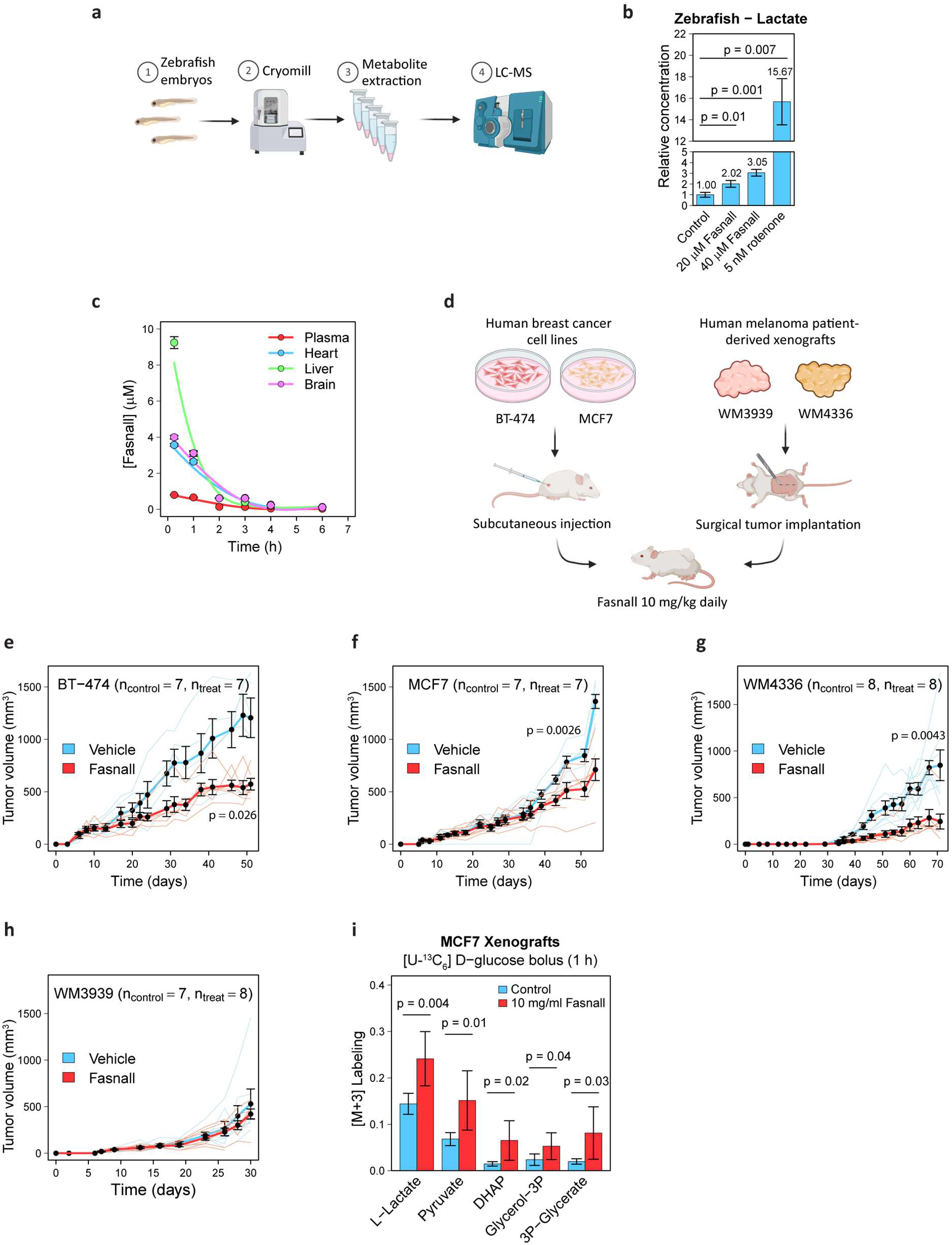
Fasnall activity *in vivo* in zebrafish embryos and mouse xenograft cancer models. **a**, Schematic of zebrafish embryo experiment. **c**, Fasnall pharmacodynamics in NSG mice after a single 10 mg/kg IP injection. **d**, Schematic of mouse xenograft experiments. **e-h**, Growth of murine tumor xenografts. i, Relative abundance of M+3 isotopologue in MCF7 tumor metabolites 1 h after IP administration of 1 g/kg [U-^13^C_6_] D-glucose bolus. Data on panels **b-c** are mean ± SD (n = 3). Data on panels **e-i** are mean ± SE (n ≥ 7). Two-sided *t* test with unequal variance was applied to the last time point.

### Fasnall pharmacokinetics in mice

To predict whether the results obtained with cancer cell lines can be translated to organismal level in mammals, we tested the pharmacokinetics of Fasnall in immunodeficient NOD*-scid* IL2Rγ^null^ (NSG) mice. In 15 min after an intraperitoneal (IP) injection of 10 mg/kg Fasnall in 50% dimethylsulfoxide/phosphate buffer saline (DMSO/PBS), Fasnall concentrations reach ∼9.2 µM in the liver, 3.5-4 µM in the heart and brain, and ∼0.8 µM in plasma, with half-life t_1/2_ 48 min, 85 min, and 88 min, respectively (Fig. 8c). DMSO was detected in all collected samples after the injection (Extended Data Fig. 9b). Measured concentrations, together with the zebrafish data, both support the premise that Fasnall has the potency to inhibit Complex I *in vivo*.

### Fasnall is effective against human cell-line-derived breast cancer and melanoma patient-derived xenografts in mice

To test whether Fasnall can inhibit tumor growth *in vivo*, we used two cancer models: (i) subcutaneous xenografts of BT-474 (HER2-enriched, ER-positive) and MCF7 (luminal B, ER-positive, derived from metastasis) breast cancer cell lines, both representing poor prognostic morphological subtypes^26,27^, and (ii) BRAF-inhibitor/MEK-inhibitor combination therapy-resistant melanoma patient-derived xenografts (PDX) WM3939 and WM4336 bearing BRAF V600E mutation. *In vitro* data from the present study have supported the sensitivity of breast cancer cells to Fasnall treatment. Combination therapy-resistant melanoma was previously shown to undergo metabolic reprogramming characterized by increased OXPHOS^28^.

Considering the fast clearance of Fasnall, we administered 10 mg/kg Fasnall via daily IP injections, starting from the day when at least half of the mice had palpable tumors (Fig. 8d). Three models responded to the treatment with a significant tumor growth decrease compared to control. One melanoma PDX model, WM3939, showed no significant difference after ∼4 weeks of treatment (Fig. 8e-h). In the models with significant response, treatment lasted ≥45 days. No significant drop in body weight was observed in mice following long-term Fasnall administration (Extended Data Fig. 9c-f). Non-steady-state labeling of lactate 1 h after administering an IP bolus of 1 g/kg [U-^13^C_6_] D-glucose with the drug or vehicle in MCF7-tumor-bearing mice demonstrated an increased lactate fermentation in tumors of Fasnall-treated mice (Fig. 8i), agreeing with our *in vitro* results. With 10 mg/kg/day Fasnall administration, the drug achieves tissue concentrations sufficient to inhibit mitochondrial respiration. In sum, Fasnall significantly slows tumor growth *in vivo* in breast cancer xenografts and in one model of combination therapy-resistant melanoma PDX.

## Discussion

Our work provides evidence of a mechanistic link between the electron transport chain (ETC) inhibition and the disruption of *de novo* fatty acid biosynthesis. Our conclusions agree with the recent finding of fatty acid biosynthesis reliance on NAD^+^ regeneration^18^ and the uncoupled demand of proliferating cells for ATP and electron acceptors^29^.

Our study shows that FASN inhibition creates multiple perturbations in the polar metabolome. We find that the minimal robust set of measurements sufficient to detect FASN inhibition in cells comprises a dose-dependent increase of the intracellular concentrations of malonate, succinate, and their CoA conjugates. While the concentrations of coenzyme A derivatives are bound by the total intracellular pool of coenzyme A, free malonate and succinate can accumulate to concentrations exceeding physiological levels by several hundredfold. Aligned with a simple substrate accumulation model, genetic silencing of FASN also increases malonyl-CoA/CoA ratios^30^, supporting our data on the pharmacological inhibition of FASN. Of course, metabolite profiling for assessing FASN activity is only valid if the compound of interest enters cells to interact with the target enzyme. For instance, GSK837149A is inactive in cell-based assays^31^ because of its inability to cross plasma membrane^20^.

An additional result of the study is that the redistribution of metabolic flux between glucose and glutamine as sources of carbon for fatty acid biosynthesis can result in an approximately unchanged sum. Therefore, isotopic tracing experiments should account for all three flux variables contributing to the fatty acid pool: (i) exogenous fatty acid consumption, (ii) *de novo* biosynthesis from glucose, and (iii) glutamine. Relying only on glucose tracing implies that glucose and glutamine fluxes are correlated for a selected treatment, and it needs to be proven first. The results of the Fasnall treatment showed that using an incomplete set of isotopic tracers can be misleading. Moreover, given that most lipid molecules are composite, the causation inference based on perturbations in the lipidome is less straightforward than using the polar metabolome. Monitoring the abundance of lipid molecules would require accounting for fluxes in a complex metabolic network spanning all lipid building blocks, making lipidome-based observations a poor surrogate for FASN inhibition. Elongase activity and exogenous fatty acid uptake are additional factors that can further complicate the readout.

Three compounds tested in the present study (Fasnall, C75, and cerulenin) did not produce the anticipated metabolic signature of FASN inhibition in cells despite being present in the cytosol. The list of doubtful FASN inhibitors can be further extended to include triclosan^32^. Previous work demonstrated that triclosan did not affect the malonyl-CoA pool at concentrations up to ∼30 µM^20^. While we cannot rule out potential perturbations in lipidome occurring via unidentified mechanisms, we conclude that Fasnall, C75, cerulenin, and triclosan are unlikely to be selective FASN inhibitors in cells and likely to have alternative enzymes as their primary targets. Such non-selective effects are not unprecedented. For instance, orlistat was reported to trigger cytidine 5’-diphosphocholine (CDP-choline) accumulation ahead of malonyl-CoA response^20^. Effects of cerulenin and orlistat on mitochondrial function were also reported previously^33^.

We classify Fasnall as a ubiquinone-dependent respiratory Complex I inhibitor similar to rotenone and metformin. The Fasnall-induced perturbations in the lipidome follow the inhibition of the TCA cycle. Like Fasnall, metformin was shown to decrease palmitate content in cancer cells^34^. We corroborate previously shown anti-cancer Fasnall activity^17,35–37^ by demonstrating its efficacy *in vivo* in breast cancer xenografts and one model of combination therapy-resistant melanoma PDX. Fasnall is not the only pharmacological agent that was found to exhibit antirespiratory activity after its introduction: recently, an ERBB2 inhibitor, ibrutinib (TAK-165), was re-assigned as a Complex I inhibitor^38,39^. Evidence for atypical Fasnall activity was published before our work^40^, where Fasnall cell toxicity was observed in FASN-knockout cells. The cell stress caused by Fasnall was found to be reproduced by arsenite, a known TCA cycle inhibitor^40^. Indeed, our work shows that FASN inhibition in cancer cells provided with exogenous sources of lipids does not strongly affect cell proliferation. However, Fasnall demonstrates a significant antiproliferative activity even in the presence of 10% dialyzed FBS.

Importantly, Fasnall-treated mice did not exhibit the side effects reported for some other Complex I inhibitors. Specifically, mice and patients treated with an OXPHOS inhibitor IACS-010759 experienced painful peripheral neuropathies such as myalgia and allodynia^4^. Unlike a previous report^17^, we observe that a 10 mg/kg IP dose of Fasnall induces transient lethargy in mice that lasts ∼30 min. This side effect was confirmed for Fasnall obtained from two different vendors (see Methods), with an MS/MS compound validation and *in vitro* metabolic tests in BT-474 cells performed for every compound batch. During lethargy and after waking up, mice do not appear to exhibit stress or pain symptoms. The fact that Complex I inhibitors do not agree in the spectrum of side effects suggests that novel selective OXPHOS inhibitors can be designed to improve the therapeutic window and minimize off-target effects. We speculate that the spectrum of side effects can also be affected by the pharmacokinetic properties of a compound. Yet, it is unclear at this time whether the Fasnall side effects will be well-tolerated by human patients.

For the FASN-specific pharmacological inhibitors, we show a previously underappreciated effect related to malonate accumulation and SDH inhibition. The consequent accumulation of succinate may have pro-tumorigenic effects^41,42^. Moreover, FASN inhibition alleviates the oxidative stress in cancer cells caused by anchorage-independent growth, which may play a role in the metastatic process^43^. To our knowledge, succinate accumulation was not previously shown for FASN inhibitors and was not reported in clinical studies.

The present work highlights the significance of reductive stress in multiple cancers. Aside from downregulating fatty acid biosynthesis, reductive stress may efficiently prevent cancer progression through multiple effects on central carbon metabolism^44,45^. The apparent NADH shuttle saturation in untreated cancer cells highlights the importance of the electron sink for cancer growth^46^. Fasnall emerges as a novel ubiquinone-dependent Complex I inhibitor available for studying mitochondrial metabolism *in vitro* and *in vivo*. In sum, NAD^+^-dependence of proliferating cancer cells creates an avenue for selective inhibitors of OXPHOS that can strengthen the arsenal of anti-cancer therapy.

### Limitations of the study

In our study, we did not assess FASN activity in a reconstituted system. Non-saponified lipids of cells treated with the panel of pharmacological agents were not profiled. The exact mechanism of Complex I inhibition by Fasnall was not pursued. The activity of Fasnall against elongases and the mechanism of cholesterol biosynthesis inhibition by Fasnall fell outside the scope of the study. While we have confirmed penetration of the pharmacological agents into BT-474 cells, the inhibitory concentrations (IC_50_) can be affected by the ability of this specific cell line to expel and transform the drugs. We have not performed metabolic flux analysis to exclude the possibility that [U-^13^C_5_] L-glutamine-derived labeling reflects an isotopic labeling exchange artifact instead of the net increase of reductive carboxylation flux^47^. However, NADH accumulation upon Fasnall treatment suggests that the law of mass action may drive the net flux in the reductive direction. The neurological side effects of Fasnall in mice were not assessed in standardized tests.

## Methods

### Key Resources

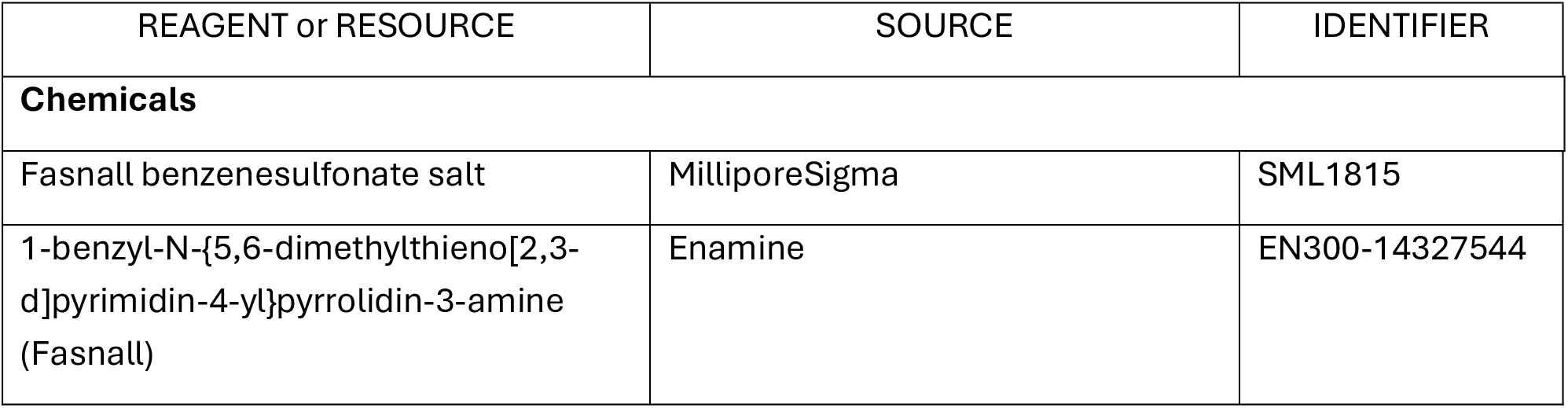
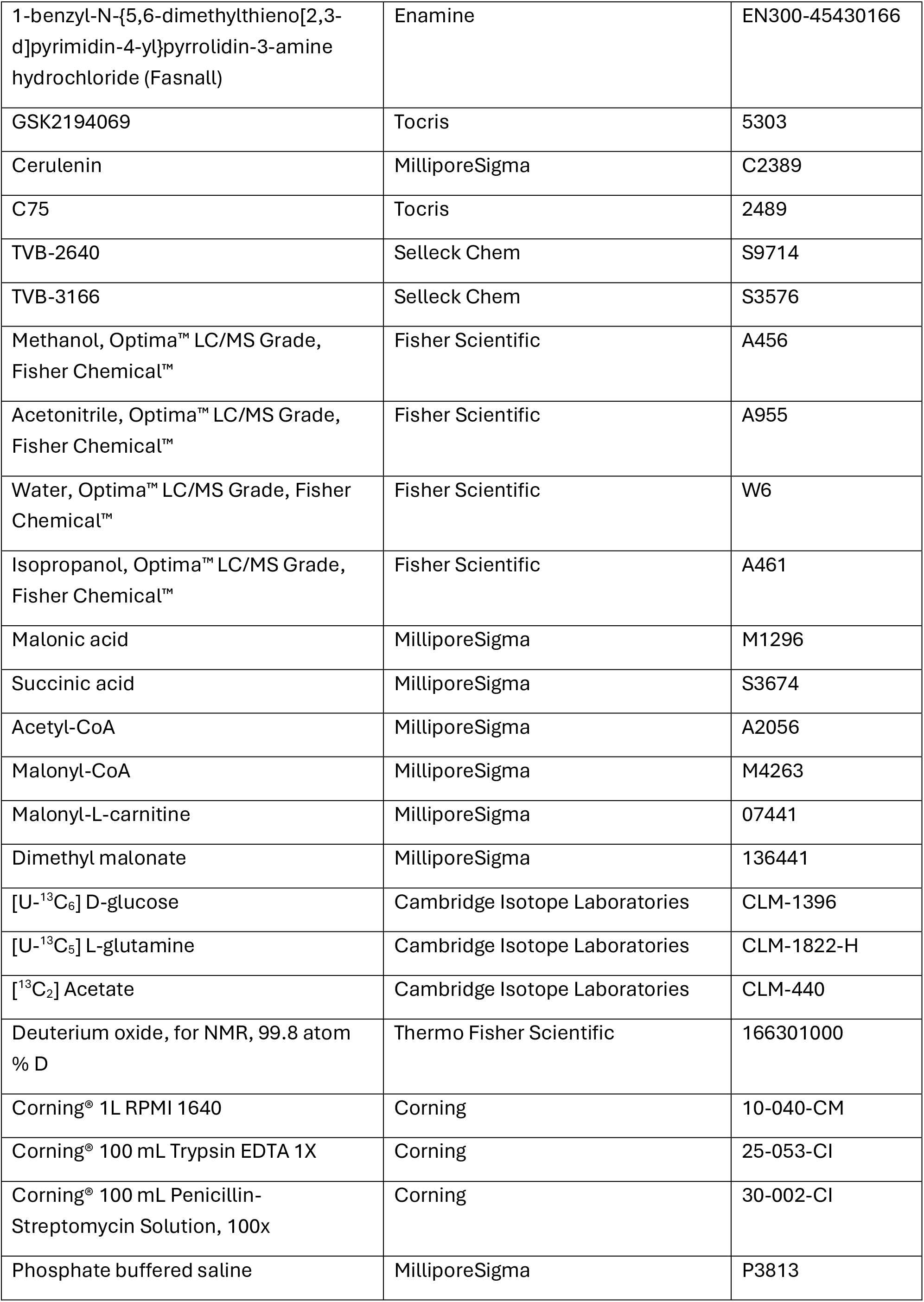
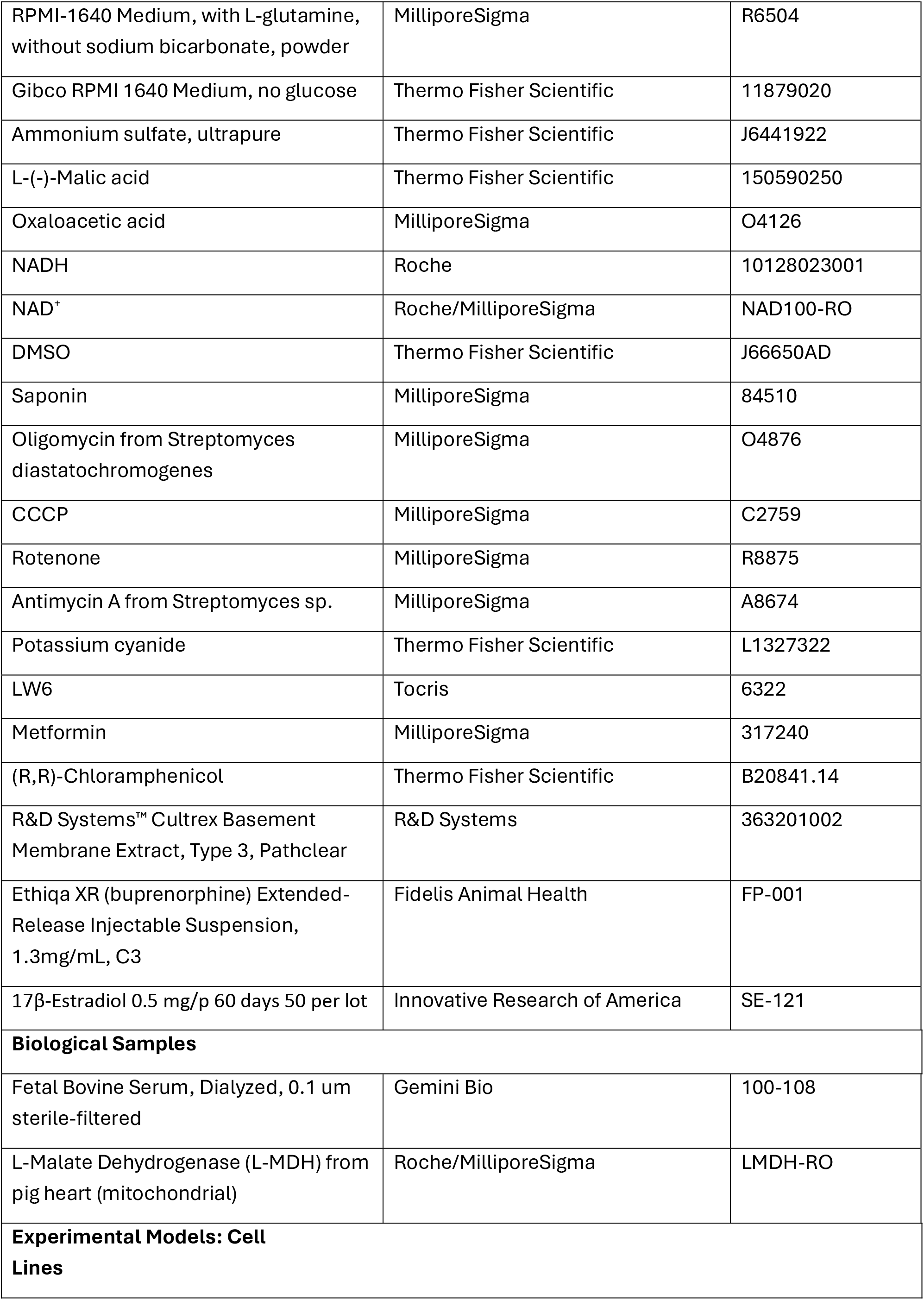
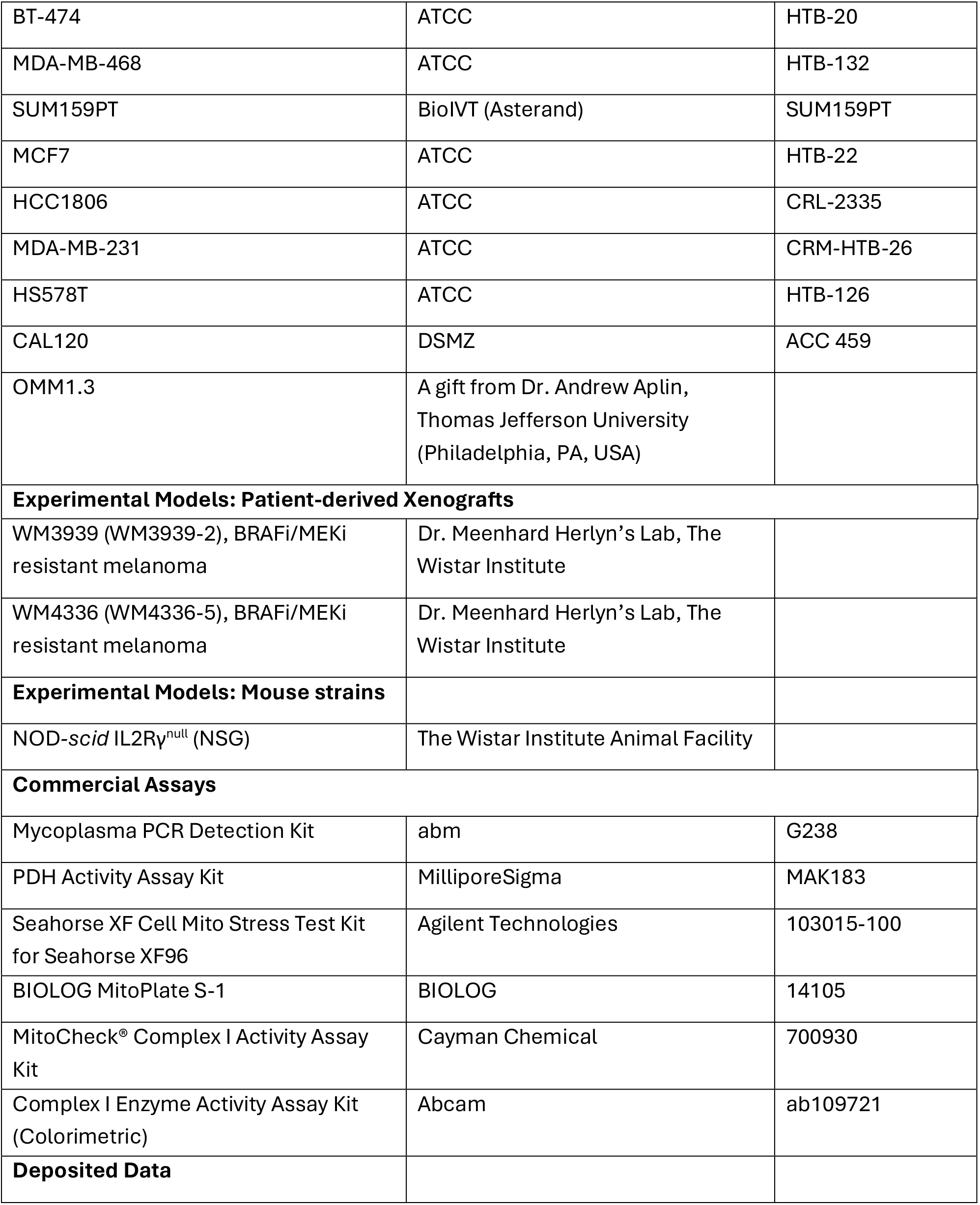
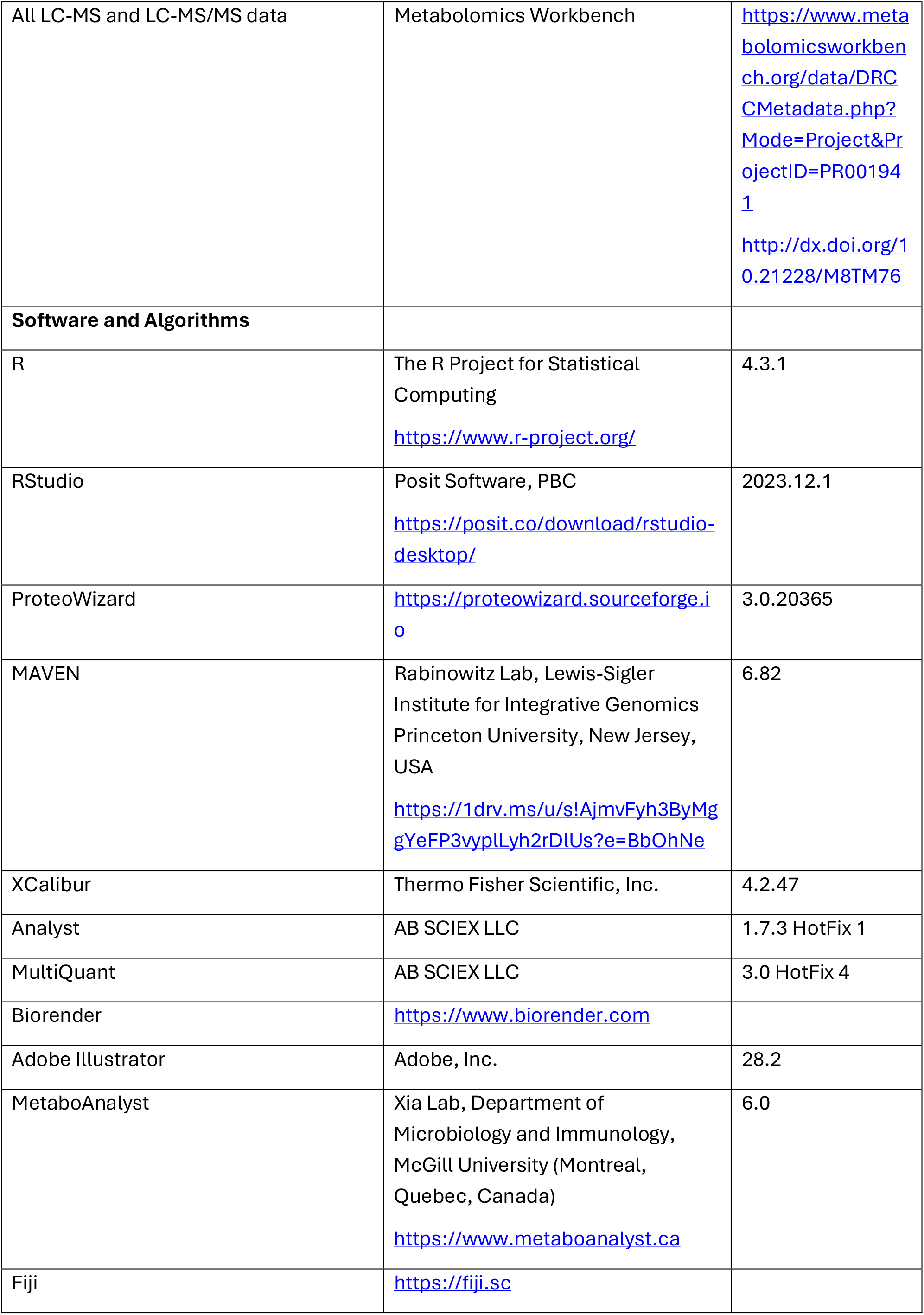

#### Cell culture

Cells were cultured in RPMI-1640 with 10% dialyzed fetal bovine serum and antibiotics. Cells were maintained in a humidified CO_2_ incubator with 37 °C and 5% CO_2_ and passaged every 2-3 days. Cell lines were regularly checked for *Mycoplasma spp.* contamination with the abm™ Mycoplasma PCR Detection Kit (cat. no. G238). For hypoxia, cells were cultured at 1% oxygen in a Coy O_2_ Control InVitro Glove Box (Coy Laboratory Products).

#### Cell proliferation assay

Cells were plated in 6-cm dishes (5·10^5^ cells per dish, 3 ml of medium) one day before the first counting. The medium was replaced at the first count with fresh medium supplemented with the corresponding drug concentration. The control condition was supplemented with 0.2% v/v DMSO. Cells were counted with Thermo Countess II FL. In brief, cells were trypsinized with 0.5 ml of trypsin. After 5 min incubation, 0.5 ml of complete medium was added to quench trypsin. Cells were resuspended by pipetting and transferred to Eppendorf tubes. After vortexing, 50 µl of cell suspension were mixed with trypan blue solution. Three replicates (separate dishes) were used per time point, one Countess chip (two chambers) per dish. Only live cells were reported. Dead cell fraction did not exceed 20% of total cells throughout all conditions and cell lines.

#### Resazurin reduction assay

Resazurin reduction was used to assess the short-term effects of drug treatment. Sterile 0.15 mg/ml resazurin stock was prepared in PBS. Cells were seeded in 96-well plates in RPMI-1640 at least 24 hours before the experiment. The medium was replaced with fresh RPMI-1640 mixed with resazurin stock 1:6 and supplemented with a drug or vehicle control. Cells were incubated for 90 min, and resorufin fluorescence was measured (560 nm excitation and 590 nm emission). The resazurin reduction assay was not used as a proxy for cell count in the present study.

#### Stable isotope tracing

For isotope tracing experiments, cells were seeded 12-16 h before the experiment. At the beginning of the experiment, the unlabeled medium was aspirated and replaced with the medium containing the isotope tracer. If not specified, the tracer concentrations matched the medium formulation for the corresponding unlabeled nutrient. For the deuterated water tracing, RPMI-1640 powder (MilliporeSigma R6504) was reconstituted in 20% D_2_O in water. Sodium bicarbonate was added according to RPMI-1640 formulation, pH was adjusted to 7.4, and the medium was sterile-filtered with a 0.2-µm filter. The experiment was conducted for a specified amount of time, followed by metabolite extraction.

#### Sample preparation for LC-MS metabolomics

For intracellular metabolite samples, the medium was aspirated, and cells were washed with PBS volume matching the volume of the medium. Metabolites were extracted with ice-cold 80% methanol. The volume of the solvent was 500 µl per 6-cm Petri dish (scaled according to the ratio of surface areas for other cell containers). After adding the methanol solution, cells were scraped from the plates, and all the content was transferred to Eppendorf tubes. For medium metabolites, 100 µl of medium was mixed with 400 µl of 100% methanol (80% final). Zebrafish embryos were combined by eight per Eppendorf tube, washed with PBS twice, and snap-frozen on dry ice. Frozen samples were ground at the temperature of liquid nitrogen by Retsch Cryomill. To each tube, 300 µl of 80% methanol were added. Plasma metabolomics samples were prepared from whole blood mixed with EDTA to 10 mM final concentration. To separate plasma, blood samples were centrifuged at 2,000 g and 4 °C for 10 min. Then, 50 µl of plasma were mixed with 450 µl of 100% methanol to reach 90% final methanol concentration. Tissue metabolite extracts were prepared from ∼50 mg snap-frozen tissue samples. Each sample was weighed while frozen. Frozen samples were ground at the temperature of liquid nitrogen by Retch Cryomill. Then, 80% methanol was added to each tube at a ratio of 1 ml per 50 mg of tissue. Regardless of the type of metabolite extracts, all samples were centrifuged at 18,000 g and 4 °C for 20 min. The supernatant was then transferred to new Eppendorf tubes and centrifuged again with the same parameters. After centrifugation, the extracts were transferred to glass LC-MS vials.

#### LC-MS metabolomics on Thermo Q Exactive HF-X

Analytes were separated on SeQuant® ZIC®-pHILIC 150 x 2.1 mm column with 5 µm particles (MilliporeSigma cat. no. 1504600001 and 1504380001). Liquid chromatography parameters were as follows. Solvent A was water with 0.01% ammonium hydroxide and 20 mM ammonium bicarbonate, solvent B – acetonitrile. A linear solvent gradient of a total duration 22.5 min was starting with 0.2 ml/min flow rate of 80% solvent B, 12.5 min – 30%, 15 min – 30%, 15.2 min – 80%, 20 min – 80%, 21 min – flow rate 0.3 ml/min, 22 min – flow rate 0.3 ml/min, 22.1 min – flow rate 0.2 ml/min. The autosampler temperature was maintained at 4 °C, the column was heated to 40 °C. The injection volume for medium and *in vivo* samples was 1 µl, for cell extracts – 5 µl.

The HESI ion source voltage was set to the following parameters: 3,600 V in both polarity modes, sheath gas 30, auxiliary gas 5, spare gas 0, probe heater 200 °C, capillary temperature 325 °C, S-Lens RF level 65. The mass spectrometer was set to acquire data in the polarity-switching mode averaging two microscans with 60,000 resolution, automatic gain control (AGC) target 5e6, scan range 72-1080 m/z, and maximum orbitrap injection time (IT) 200 ms. Data were converted into the mzXML format by ProteoWizard and analyzed in MAVEN. All ^13^C isotopologue abundances were deisotoped (unless stated otherwise) by considering the natural abundance of ^13^C and the isotope enrichment of the tracers.

#### LC-MS metabolomics on SCIEX QTRAP5500

The chromatography method on the SCIEX LC-MS system was identical to the one described above. ESI source parameters were as follows: curtain gas 35, collision gas “Medium”, ion spray voltage 4500 V for both polarity modes, probe temperature 500 °C, ion source gas 1 and 2 at 70. Data were recorded in the polarity-switching mode with scheduled advanced multiple-reaction monitoring (MRM) with most peaks acquired in 120 s windows. Pause between mass ranges 2 ms, minimum dwell time 2 ms, maximum – 250 ms. MRM parameters were optimized by the direct injection of pure chemical standards. Details of the MRM tables are available in the deposited mass spectrometry data. Data were analyzed in MultiQuant.

#### Sample preparation for LC-MS free fatty acid analysis

Cells were seeded in 10 cm Petri dishes. At collection, cells were washed with PBS three times. Then, 1 ml of methanol was added, and cells were scraped from the surface. All content of the plate was transferred into 13 x 100 mm Pyrex glass tubes. Lipids were extracted by the Folch method. Dried lipids were redissolved in 1 ml of 0.3 M KOH solution in 90% methanol and incubated at 85 °C for 1 h. Then, 100 µl of formic acid were added, followed by 800 µl of hexane for extraction. The hexane phase was transferred to glass LC-MS vials and dried under the stream of nitrogen. Samples were redissolved in 1 ml of 1:1 methanol:isopropanol.

#### LC-MS free fatty acid analysis on Thermo Q Exactive HF-X

Analytes were separated on a Phenomenex Kinetex XB-C18 150 x 3 mm column, 2.6 µm particle size, 100 Å pore size (cat. no. 00F-4496-Y0, AJ0-8775, and AJ0-9000). Liquid chromatography parameters were as follows. Solvent A was 60:40 acetonitrile:water with 10 mM ammonium formate, solvent B – 90:8:2 isopropanol:acetonitrile:water with 10 mM ammonium formate. Mobile phase composition was changing according to the following liner gradient program (with respect to solvent B): 0 min – 15% and flow rate 0.333 ml/min; 4.5 min – 60%; 12 min – 82%; 12.75 min – 95%; 16.5 min – 100%, 22.5 min – 100%; 22.6 min – 15% and flow rate 0.5 ml/min, 24.9 min – 15% and flow rate 0.5 ml/min, 25 min – 15% and 0.333 ml/min. The autosampler temperature was maintained at 20 °C, the column was heated to 65 °C.

The HESI ion source voltage was set to the following parameters: +3,300/-3,500 V, sheath gas 50, auxiliary gas 20, spare gas 0, probe heater 350 °C, capillary temperature 350 °C, S-Lens RF level 80. The mass spectrometer was set to acquire data in polarity-switching mode, one microscan, 60,000 resolution, AGC target 5e6, scan range 130-1950 m/z, IT 200 ms. Data were converted into the mzXML format by ProteoWizard and analyzed in MAVEN. Free cholesterol was measured as a water-loss in-source fragment [M-H_2_O+H]^+^.

#### Drug MS^2^ CID profiles on SCIEX QTRAP5500

GSK2194069 and Fasnall CID profiles were recorded for 10 µl/min direct injection in 50% pHILIC solvent B in acetonitrile in the positive mode with the ion source parameters described above for the metabolomics method. The concentrations were 50 ng/ml for GSK2194069 and 1 ng/ml for Fasnall. Spectra were acquired in the Enhanced Product Ion (EPI) mode using the third quadrupole as an ion trap, with unit resolution, declustering potential (DP) 50 V, entrance potential (EP) 10 V, settling time 100 ms, and dynamic fill time. Collision energy varied from 5 to 55 V with a 5 V step; each spectrum was produced by averaging 4 scans.

#### Pathway analysis

Pathway enrichment analysis was performed in MetaboAnalyst 6.0 (www.metaboanalyst.ca) using the list of metabolites with significantly differentiated abundance (Supplementary Table 2). Topology analysis was set to “Out-degree Centrality”; the pathway library – “Homo sapiens (SMPDB).”

#### Seahorse assay

Oxygen consumption rate (OCR) was measured on a Seahorse XFe96 extracellular flux analyzer (Agilent Technologies) following the manufacturer’s instructions. In brief, BT-474 cells were plated without inhibitors in DMEM/F12 12 h before the experiment, 50,000 cells per well. During the assay, after recording the OCR baseline, the following compounds were injected sequentially (final concentrations provided): 5 µM GSK2194069 or Fasnall, 2.5 µM oligomycin, 5 µM CCCP, and 5 µM rotenone together with 20 µM antimycin A.

#### BIOLOG mitochondrial function assay

The assay mix was prepared according to the manufacturer’s instructions with the saponin final concentration 50 µg/ml (1.2 mg/ml stock in sterile water). BIOLOG S-1 plates were pre-incubated with the assay mix (30 µl per well) containing twice the final concentration of the corresponding drug (Fasnall or GSK2194069) for 1 hour at 37 °C. After the pre-incubation, BT-474 cells were trypsinized, washed with PBS twice, and resuspended in BIOLOG 1x MAS buffer at the concentration 1e6 cells per ml. Then, 30 µl of cell suspension were transferred to each well (30,000 cells per well). The plate reader maintained 37° C during the experiment. Absorbance at 590 and 750 nm was recorded every 5 min for 5 hours. Long-wavelength absorbance was used as a baseline to correct the artifacts of measurements. The initial linear part of the kinetic profile was used to estimate the reaction rate via linear regression. Rates were measured in triplicates. Welch’s t-tests, followed by the Benjamini-Hochberg false discovery rate control, were used to estimate the statistical significance.

#### PDH enzymatic assay

36 million BT-474 cells were lysed in 3.6 ml of the PDH assay buffer in the Dounce homogenizer with pestle A. Two-thirds (2.4 ml) were mixed with saturated ammonium sulfate (NH_4_)_2_SO_4_ (∼4.1 M) in proportion 1:2 to precipitate proteins. The mixture was centrifuged at 10,000 g for 5 min at 4 °C. The protein pellet was floating on the surface of the mixture. The protein pellet was immersed in a fresh ammonium sulfate solution and centrifuged again. The ammonium sulfate solution was carefully removed. The pellet was resuspended in 1.2 ml of the PDH assay buffer. The PDH reaction was conducted according to the manufacturer’s instructions.

#### MDH enzymatic assay

MDH activity was reconstituted in the assay buffer containing 40 mM Tris pH 7.4, 200 mM NaCl, and 4 mM EDTA. The final volume per reaction was 30 µl consisting of 14 µl of the 2x assay buffer, 1 µl of the 1:100 MDH dilution in the 2x assay buffer (1.67 µg/ml final concentration), 2.5 µl of 12.5 mM NADH (1.04 mM final), 2.5 µl of 12.5 mM oxaloacetic acid, and 10 µl of water. To collect samples, 5 µl of the reaction solution was mixed with 500 µl of 80% methanol. The samples were centrifuged once at 18,000 and 4 °C for 20 min and analyzed with the SCIEX QTRAP5500 LC-MS metabolomics method described above.

#### Zebrafish pharmacological inhibitor assays

The zebrafish research was approved by the University of Pennsylvania Institutional Animal Care and Use Committee. Husbandry was performed in accordance with institutional animal welfare guidelines. Embryos from wild type, Tübingen zebrafish were collected within 15 minutes of fertilization. Embryos between 2 and 3 crosses were equally pooled and allocated to treatment conditions for all experiments. Embryos were reared at 28 °C in E3 medium (4 mM NaCl, 0.17 mM KCl, 0.33 mM CaCl_2_, and 0.33 mM MgSO_4_, no methylene blue) until treatment. Pharmacological agents were diluted from stocks at room temperature in E3 for 15 minutes before treatment. For treatment at 48 h post-fertilization, embryos in their chorions were transferred to clean plates with the inhibitor-supplemented E3 and incubated for 6 hours. Embryos were dechorionated and imaged in E3 using a Leica IC80HD camera. Images were processed in FIJI.

#### Fasnall pharmacodynamics in mice

18 NOD*-scid* IL2Rγ^null^ (NSG) female mice were injected intraperitoneally (IP) with 10 mg/kg of Fasnall in 1:1 DMSO:PBS (50 µl of 5.4 mg/ml Fasnall per injection). Three animals were sacrificed at each time point. Blood was collected via the intracardiac puncture, and EDTA was added to the final concentration of 10 mM immediately. Tissue samples were snap-frozen on dry ice.

#### Mouse xenograft experiments

All animal experiments were approved by the Institutional Animal Care and Use Committee at the Wistar Institute and were performed in an Association for the Assessment and Accreditation of Laboratory Animal Care-accredited facility. NSG-strain 6-10 week-old female mice were used in the study, with eight animals per group. For MCF7 and BT-474 xenografts, 2 million cells were injected subcutaneously into the flanks of the animals in 100 µl of a 1:1 mix of DMEM and Cultrex Basement Membrane Extract, Type 3. Immediately after injecting cells, 60-day release 0.5 mg 17β-estradiol pellets were inserted subcutaneously by a trocar size 10 into the interscapular area. The procedure was performed under general anesthesia with isoflurane. Starting from the day when at least half of the mice had palpable tumors, mice were treated daily by IP injections of 50 µl of 5.4 mg/ml Fasnall prepared in a 1:1 DMSO:PBS mix. This dosage corresponds to 10 mg/kg for the average animal weight in the study (∼26 g). The control groups received 50-µl injections of 1:1 DMSO:PBS. Tumors were measured by a caliper, and tumor volume was calculated as 0.5 × *L* × *W*^2^, where *L* and *W* are the length and width, correspondingly.

Patient-derived xenograft tumor tissue pieces 3×3 mm were surgically implanted subcutaneously in the lumbar area under general anesthesia with isoflurane as described elsewhere^48^. Mice received slow-release analgesia after the procedure (3.25 mg/kg Ethiqa XR). During the first two weeks after surgery, mice were treated with 0.25 mg/ml amoxicillin in drinking water.

## Statistics

All statistical tests using experimental data were performed in R. No statistical method was used to predetermine sample sizes. No data were excluded from the analyses. The experiments were not randomized, and the investigators were not blinded to allocation during experiments and outcome assessment.

## Resource availability

### Lead Contact

Further information and requests for resources and reagents should be directed to and will be fulfilled by the Lead Contact, Dr. Zachary T. Schug.

### Materials Availability

Reagents and materials used to conduct the research detailed in this manuscript are available on request from the Lead Contact, Dr. Zachary T. Schug.

### Data and Code Availability

The LC-MS data presented in the study were deposited on www.metabolomicsworkbench.org under the project identifier PR001941 and the DOI http://dx.doi.org/10.21228/M8TM76. All data are available from the corresponding author upon request.

### Supplemental Item Titles

Supplementary Table 1 | Clinical trials of FASN inhibitors in cancer patients.

Supplementary Table 2 | The lists of significantly differentiated metabolites used as queries for the pathway analysis in MetaboAnalyst 6.0.

Supplementary Table 3 | Pathway enrichment analysis results for GSK2194069-treated BT-474 cells.

Supplementary Table 4 | Pathway enrichment analysis results for Fasnall-treated BT-474 cells.

## Acknowledgments

We thank Dr. Meenhard Herlyn (The Wistar Institute) and his lab members for supporting the study and providing melanoma patient-derived xenografts, as well as Dr. Andrew Aplin (Thomas Jefferson University, Philadelphia, PA, USA) for providing OMM1.3 cell line. We thank Dr. Hsin-Yao Tang, Dr. Aaron Goldman, Nicole Gorman, and Thomas Beer of the Wistar Institute Proteomics & Metabolomic Core, which were supported, in part, by NIH grant S10 OD023586. We would like to thank the staff of the Wistar Animal Facility, Molecular Screening & Protein Expression Facility, and Humanized Models of Disease Core for their support. Many thanks to S. Luke Blair for valuable discussions. This work was financially supported by grants from the National Institutes of Health (NIH), National Cancer Institute (NCI) DP2 CA249950-01, NIH NCI P01 CA114046, Melanoma Research Foundation 717173, and Susan G. Komen CCR19608782.

## Author Contributions

D.M. and Z.T.S. conceived the study, designed the experiments, and analyzed the experimental results. D.M. and J.D. performed cell proliferation assays. D.M. and S.O. performed isotope tracing and *in vitro* enzymatic assays. S.O. measured oxygen consumption. D.M., J.D., K.P., and F.B. performed unlabeled metabolomics experiments. J.P. and M.M. designed and performed the zebrafish embryo assay. D.M. performed mass spectrometry analysis and mouse experiments. D.M. wrote the manuscript with input from all authors. All authors read and approved the manuscript.

## Declaration of Interests

Z.T.S. is a scientific co-founder and consultant for Syndeavor Therapeutics. Other authors declare no competing interests.

## Extended Data

**Extended Data Fig. 1.**
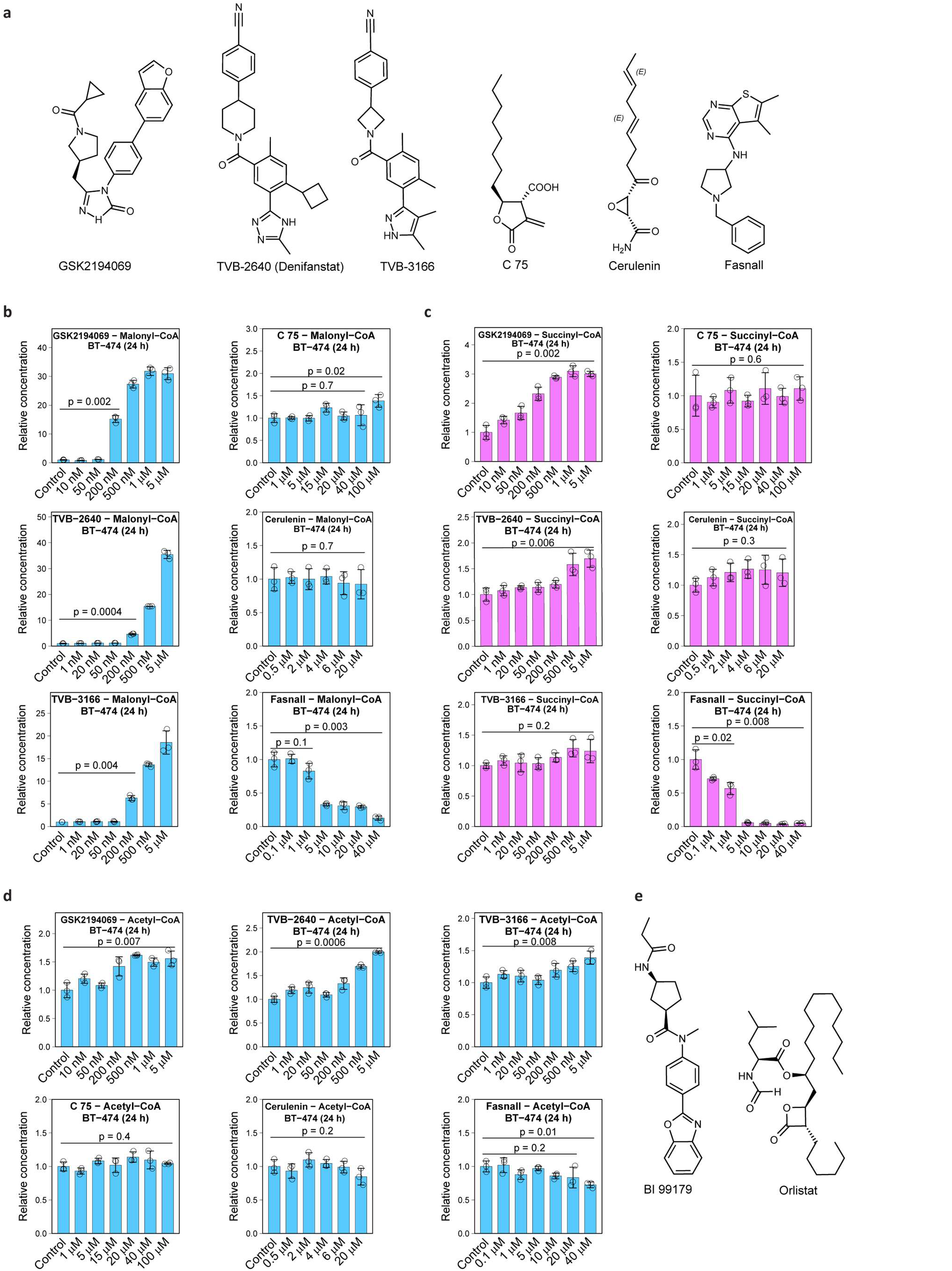
FASN inhibitors and their effect on the relative concentrations of coenzyme A species. **a**, Chemical structures of six previously described FASN inhibitors used in the present study. **b-d**, Relative concentrations of malonyl-CoA (**b**), succinyl-CoA (**c**), and acetyl-CoA (**d**) in BT-474 cells treated with six FASN inhibitors. **e**, Chemical structures of BI 99179 and orlistat. Data are mean ± SD, n = 3.

**Extended Data Fig. 2.**
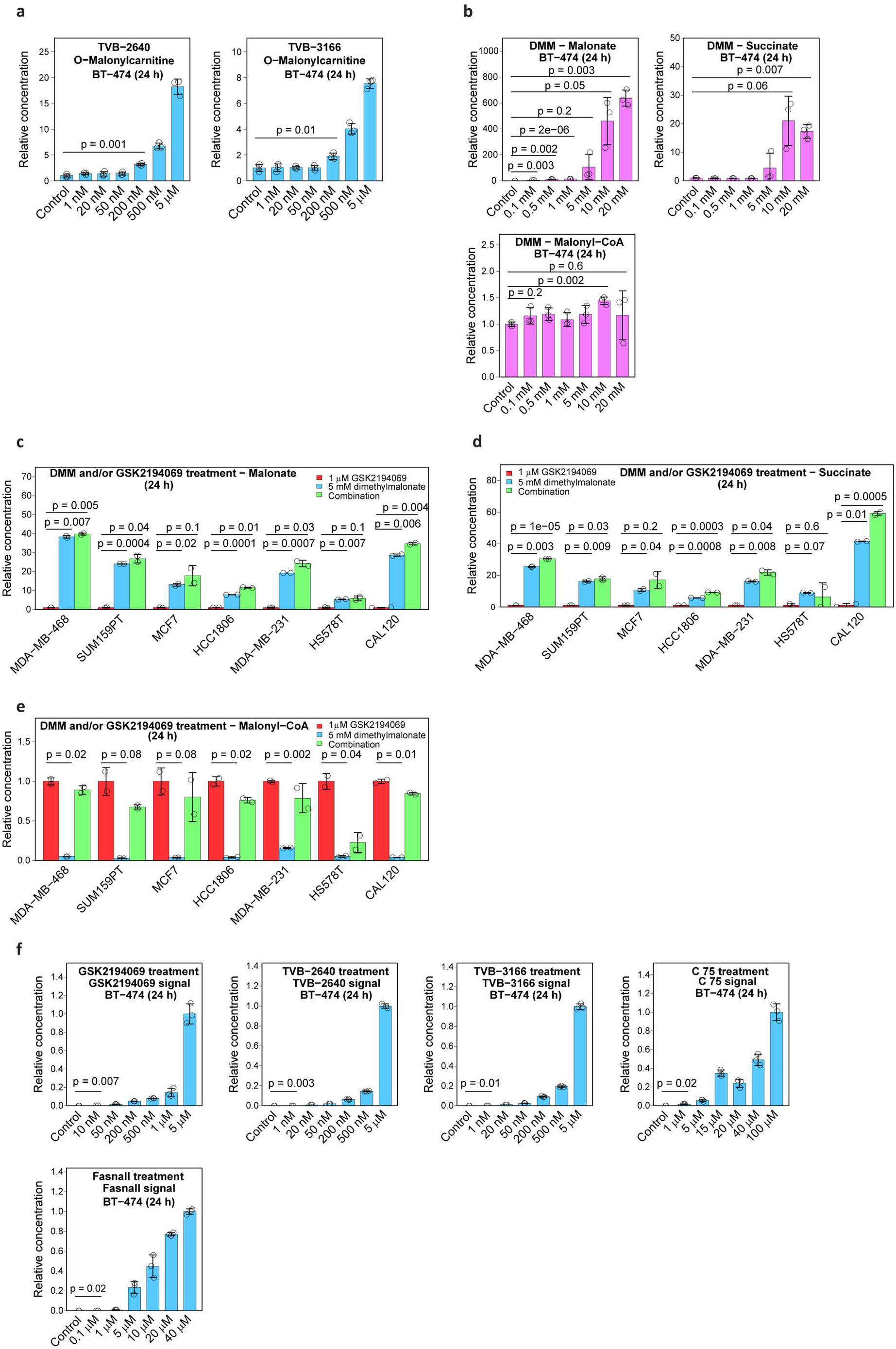
The aspects of metabolic perturbations caused by FASN inhibitors and dimethylmalonate and the assessment of their intracellular presence. **a**, O-Malonyl-L-carnitine concentrations in BT-474 cells treated with TVB-2640 and TVB-3166. **b**, Malonate, succinate, and malonyl-CoA concentrations in BT-474 cells treated with DMM. **c-e**, Malonate (**c**), succinate (**d**), and malonyl-CoA (**e**) concentrations in seven breast cancer cell lines treated with DMM and/or GSK2194069 confirming the metabolic signature of SDH inhibition by malonate. **f**, Intracellular concentrations of GSK2194069, TVB-2640, TVB-3166, C75, and Fasnall upon treatment of BT-474 cells i*n vitro*. Data are mean ± SD, n = 3.

**Extended Data Fig. 3.**
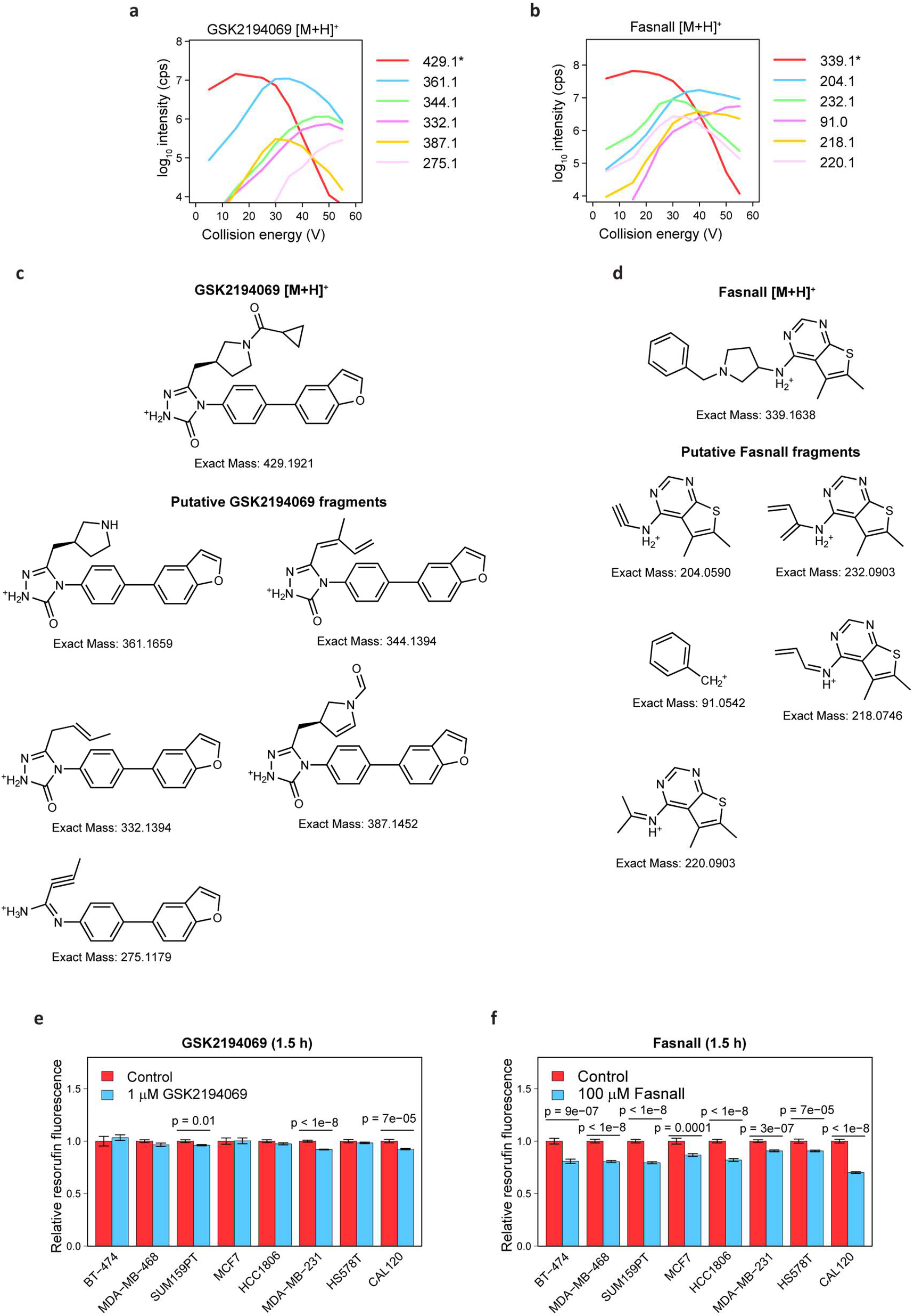
MS/MS fragmentation of GSK2194069 and Fasnall and the short-term effect of the two drugs on the resorufin-to-resazurin conversion in eight breast cancer cell lines. **a-b**, CID profiles recorded on SCIEX QTRAP5500 for GSK2194069 (**a**) and Fasnall (**b**) depicting the parental ion (red) and five major fragments. **c-d**, Putative fragment structures for GSK2194069 (**c**) and Fasnall (**d**) matching the experimentally observed ion masses. **e-f**, Relative resazurin fluorescence in eight breast cancer cell lines treated with 1 µM GSK2194069 (**e**) and 100 µM Fasnall (**f**) for 1.5 h. Data in panels **e-f** are mean ± SE, n ≥ 12.

**Extended Data Fig. 4.**
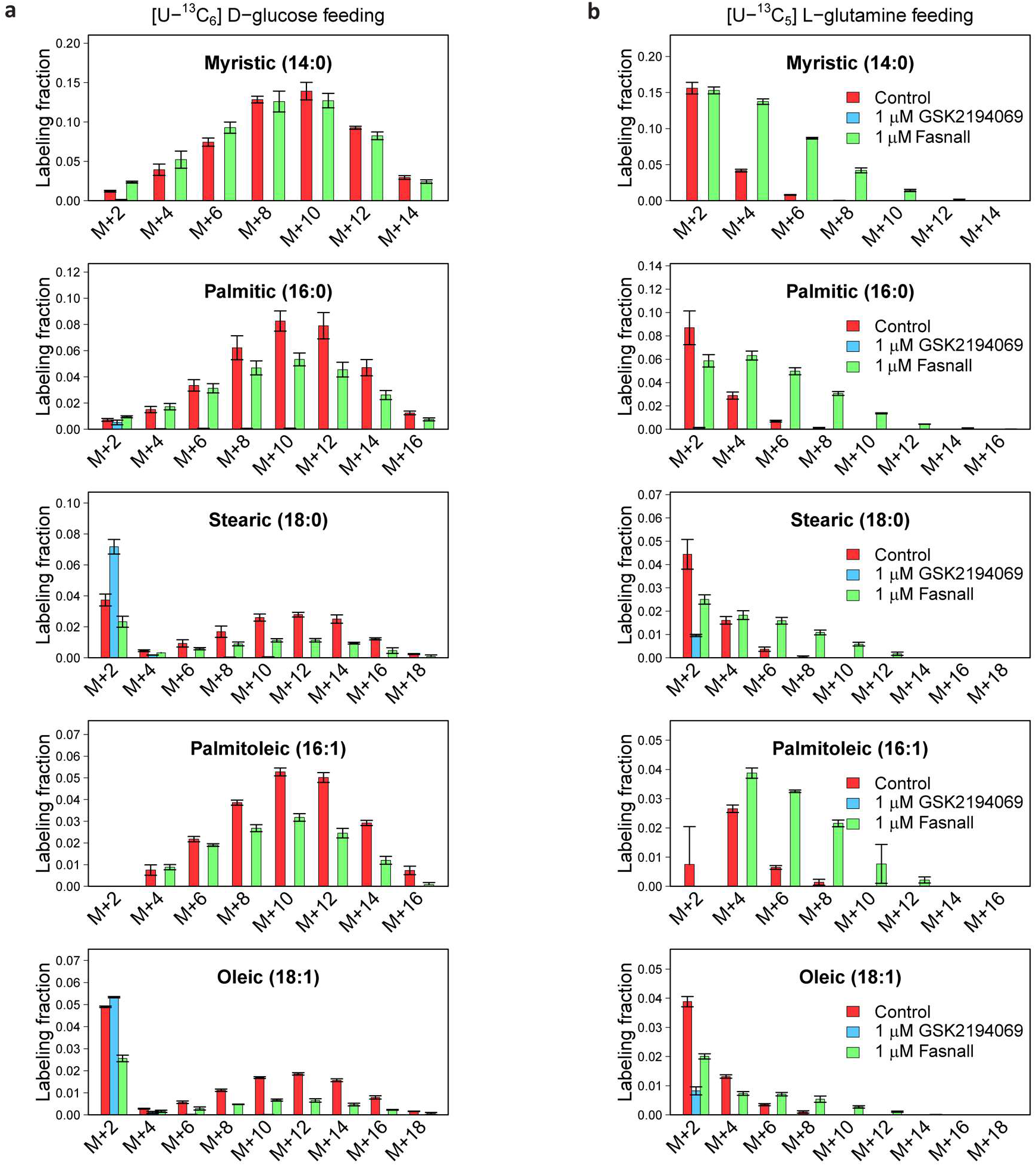
Isotopologue distributions of fatty acids in BT-474 cells treated with 1 µM GSK2194069 and 1 µM Fasnall. **a**, [U-^13^C_6_] D-glucose tracing. **b**, [U-^13^C_5_] L-glutamine tracing. Data are mean ± SD, n = 3.

**Extended Data Fig. 5.**
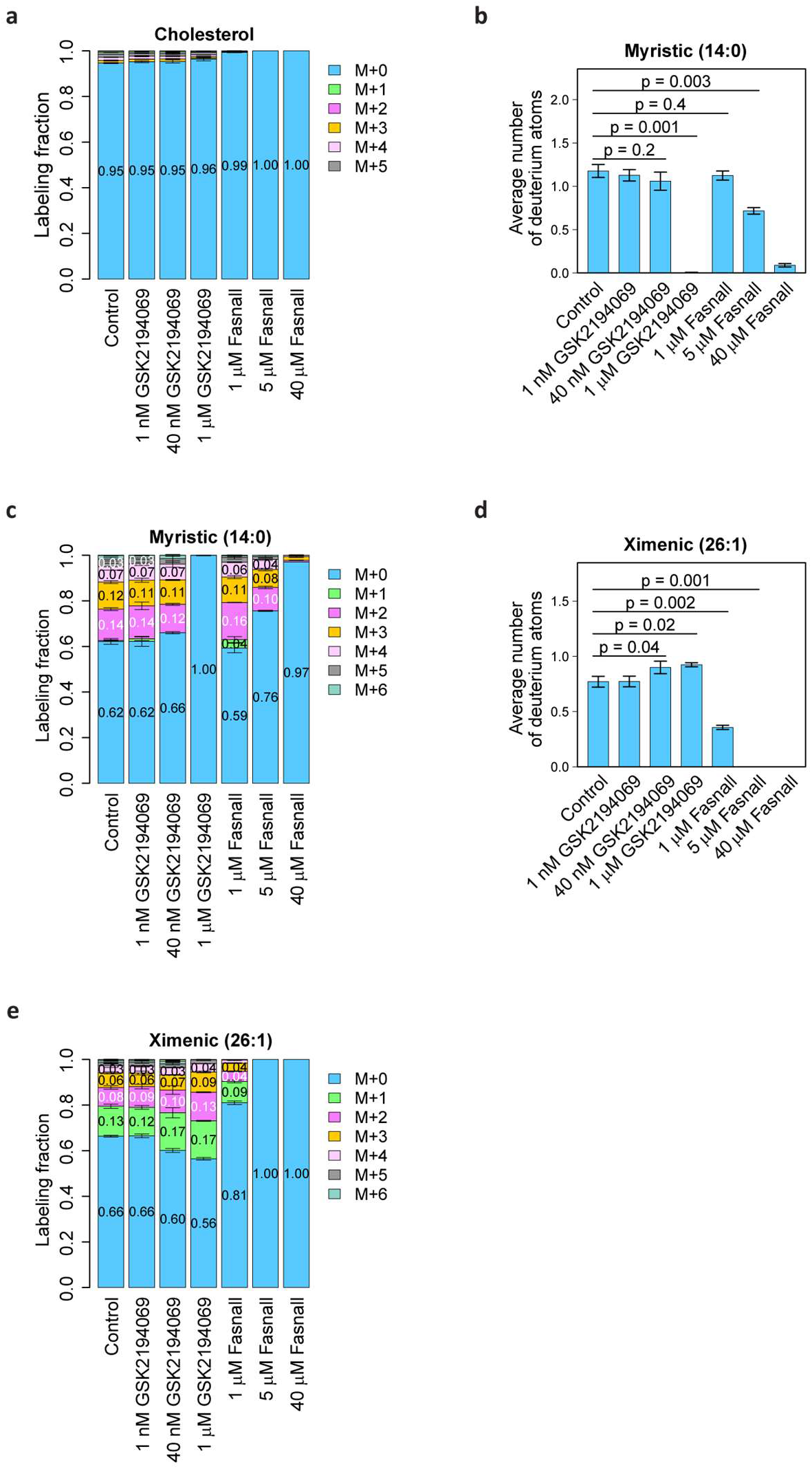
Deuterium labeling of cholesterol, as well as long-chain (myristic) and very-long-chain (ximenic) fatty acids upon treatment with GSK2194069 Fasnall in BT-474 cells grown in 20% D_2_O medium. **a**, Cholesterol deuterium isotopologue distribution. **b-c**, Total deuterium labeling (**b**) and the isotopologue distribution (**c**) for myristic acid. **d-e**, Total deuterium labeling (**d**) and the isotopologue distribution (**e**) for ximenic acid. Data are mean ± SD, n = 3.

**Extended Data Fig 6.**
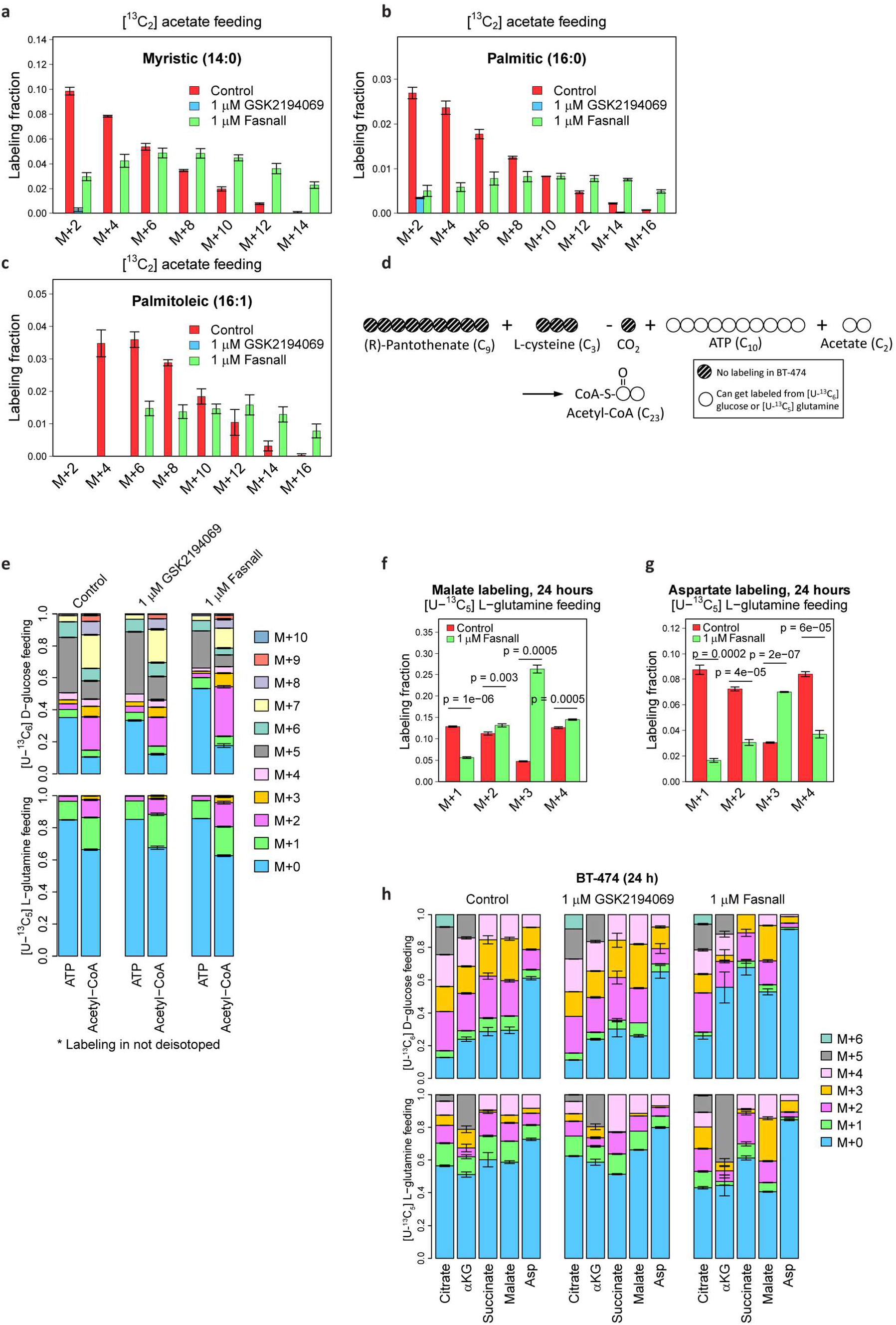
Details of the 24 h isotope tracing in BT-474 treated with 1 µM GSK2194069 and 1 µM Fasnall. **a-c**, Isotopologue distributions of myristic (**a**), palmitic (**b**), and palmitoleic (**c**) acids in cells fed with 200 µM [^13^C_2_] acetate. **d**, Schematic of acetyl-CoA carbon labeling in cells fed with [U-^13^C_6_] D-glucose or [U-^13^C_5_] L-glutamine. **e**, Metabolite labeling data used for inferring acetyl group labeling on acetyl-CoA molecule. **f-g**, Malate (**f**) and aspartate (**g**) labeling in BT-474 cells fed with [U-^13^C_6_] D-glucose and [U-^13^C_5_] L-glutamine. **h**, Complete isotopologue distributions of the TCA cycle-associated metabolites in BT-474 cells fed with [U-^13^C_6_] D-glucose and [U-^13^C_5_] L-glutamine for 24 h. Data are mean ± SD, n = 3.

**Extended Data Fig. 7.**
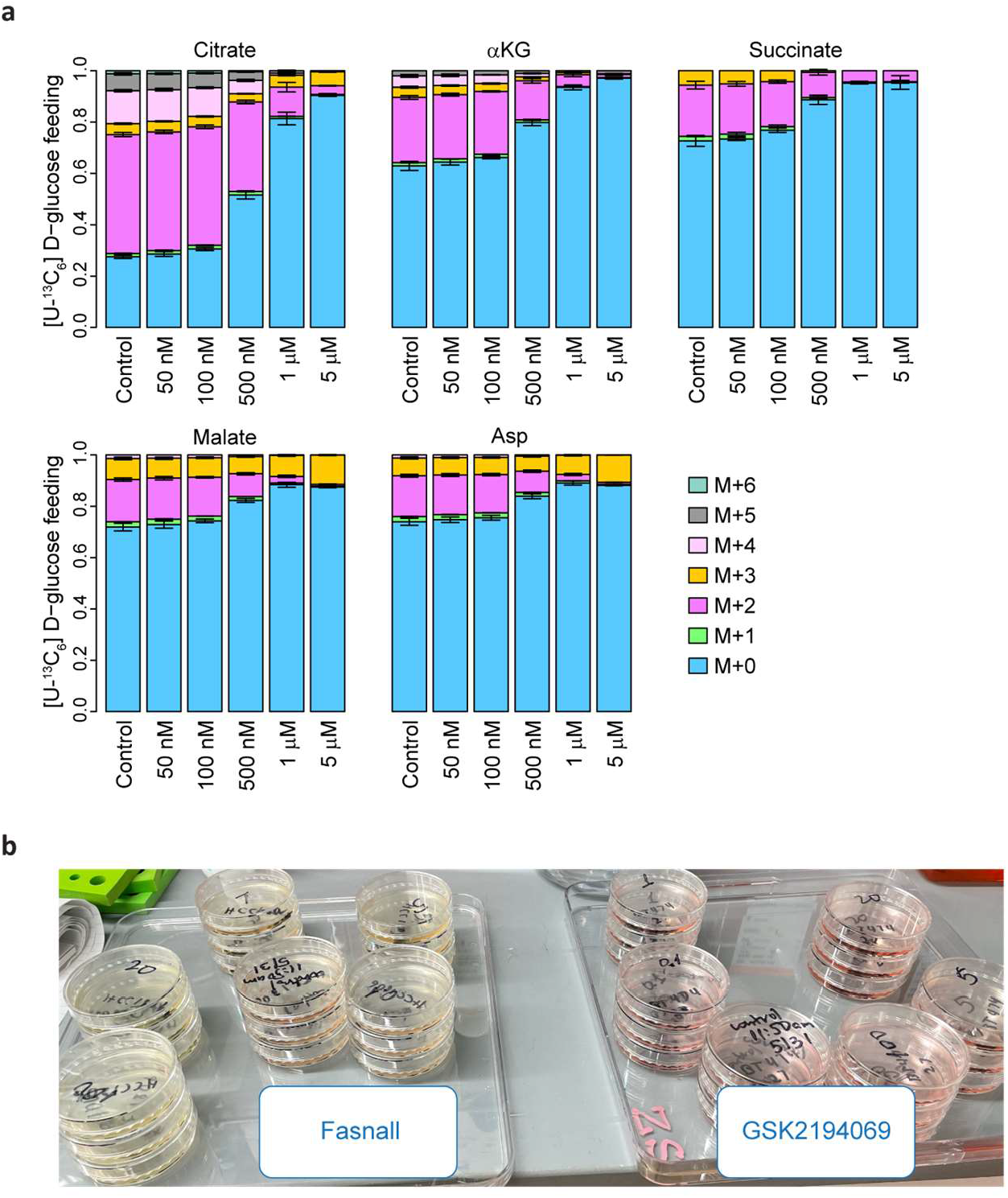
Details of the 4 h [U-^13^C_6_] D-glucose isotope tracing in cells treated with 1 µM GSK2194069 and 1 µM Fasnall. **a**, Complete isotopologue distributions of the TCA cycle-associated metabolites in OMM1.3 cells fed with [U-^13^C_6_] D-glucose for 4 h. b, Medium acidification in Petri dishes with BT-474 cells treated with Fasnall (left) or GSK2194069 (right) for 12 h. Data are mean ± SD, n = 3.

**Extended Data Fig. 8.**
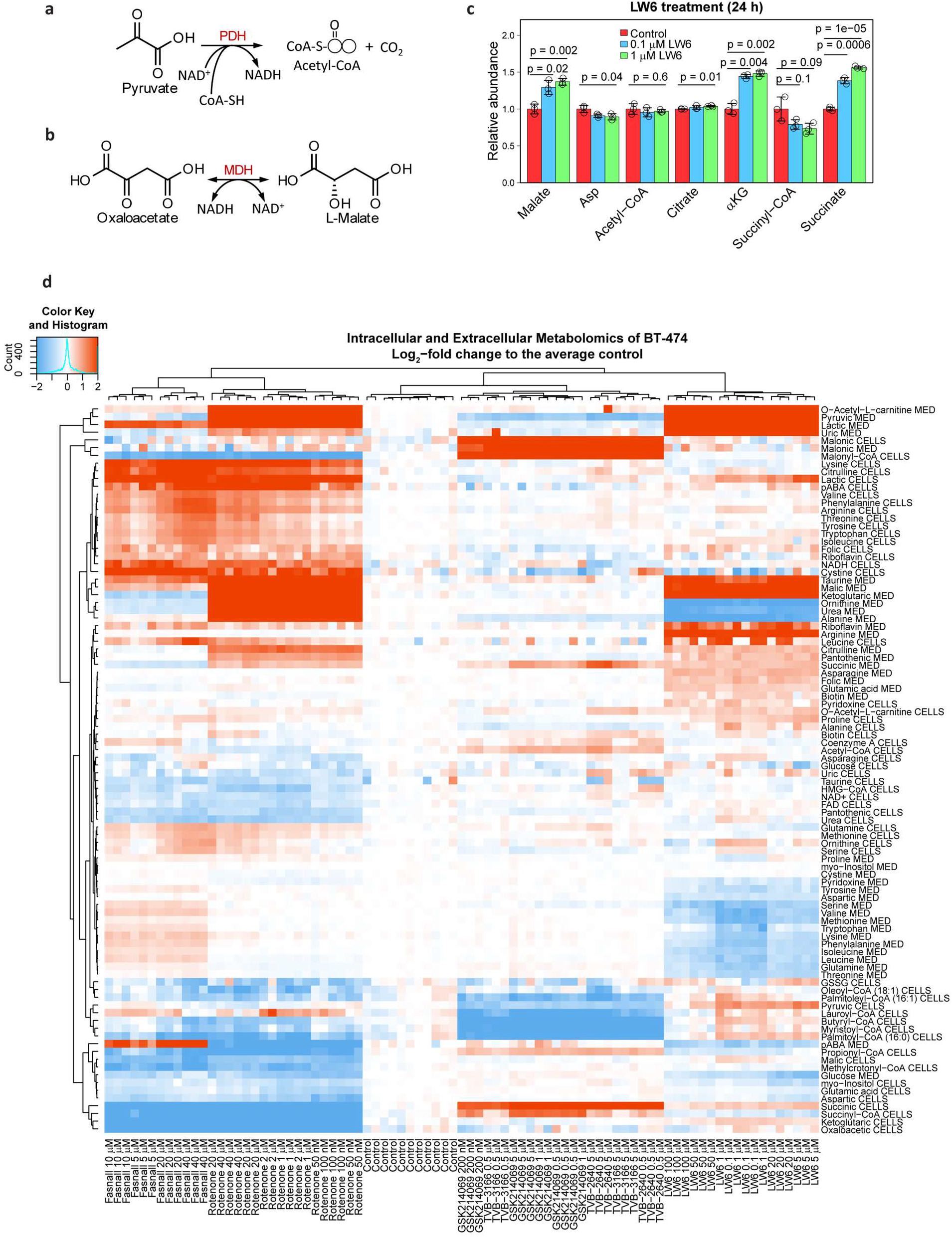
Fasnall metabolic activity matches Complex I inhibition. **a-b**, Chemical reactions monitored in PDH (**a**) and MDH (**b**) assays. **c**, The effect of 24-h LW6 treatment on metabolite abundance in BT-474 cells. Data are mean ± SD, n = 3. **d**, Relative metabolite concentration changes in BT-474 cells treated with a panel of pharmacological agents.

**Extended Data Fig. 9.**
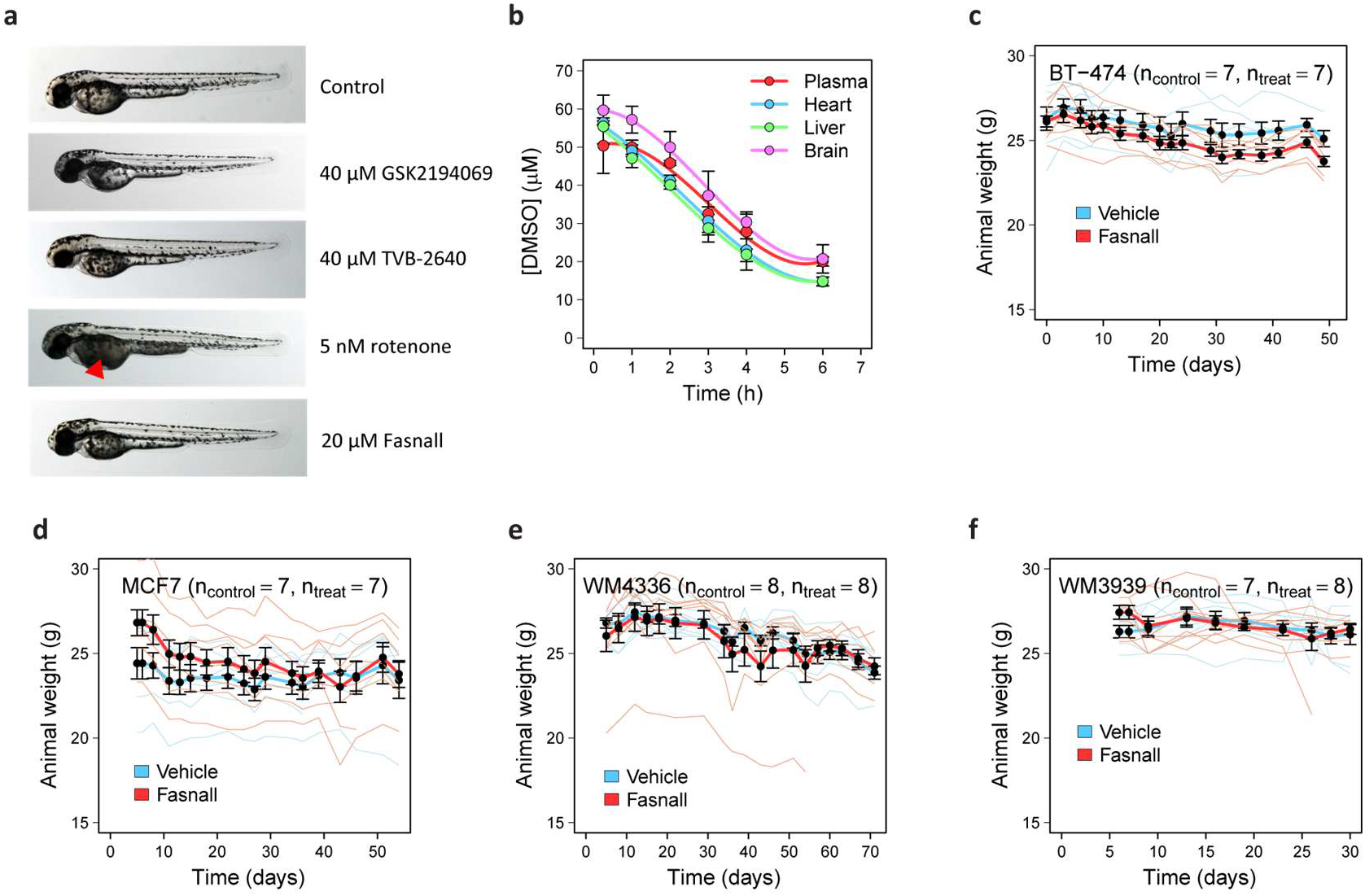
Details of in vivo experiments with Fasnall. **a**, Morphology of zebrafish embryos treated with GSK2194069, TVB-2640, rotenone, and Fasnall. **b**, DMSO (vehicle of Fasnall injection) pharmacodynamics. **c-f**, Animal weight in tumor xenograft experiments with BT-474 (**c**), MCF7 (**d**), WM4336 (**e**), and WM3939 (**f**) xenografts. Data on panel **b** are mean ± SD, n = 3. Data on panels **c-f** are mean ± SE, n ≥ 7.

## Notes

https://www.metabolomicsworkbench.org/data/DRCCMetadata.php?Mode=Project&ProjectID=PR001941

